# Androgen receptor interactions provide insight into steroid mediated metabolic shifts in endocrine resistant breast cancer

**DOI:** 10.1101/2024.11.01.621520

**Authors:** Rachel Bleach, Emir Bozkurt, Katherine Sheehan, Sally Shirran, Jingqi Xin, Stephanie Agbana, Mihaela Ola, Leonie Young, Michael W O’Reilly, Jochen HM Prehn, Marie McIlroy

## Abstract

**Purpose:** Aromatase Inhibitors (AI) are standard therapy for hormone receptor positive breast cancers in post-menopausal patients. Disease recurrence is common and previous studies suggest that the altered steroid environment may be a driver of resistance. Using label-free mass-spectrometry we explored the unique androgen receptor (AR) interactome that supervenes in AI resistant breast cancer and the associated hyperandrogenic environment.

**Experimental Design:** AR expression was evaluated in a primary breast cancer tissue-microarray (n=875) with nuclear and cytoplasmic localization quantified. Liquid-chromatography tandem mass-spectrometry (LC-MS/MS) analysis was utilized to identify proteins interacting with the AR in models of AI-resistance. Validation was carried out by co-immunoprecipitation and co-localisation studies. Live-cell imaging, Seahorse MitoStress Assays and flow cytometry were used to quantify changes in mitochondria and cell metabolism arising in models of AI-resistance.

**Results:** Utilising digital pathology we detected that abundant cytoplasmic AR protein was associated with poor survival only in the post-menopausal cohort, and most significantly, in the therapy-refractory Luminal B subtype (p=0.0085). Models of AI-resistance and androgen excess highlight diffuse AR localisation throughout the cytoplasm and nucleus accompanied by increased mitochondrial mass and membrane potential, and increased oxidative phosphorylation and glycolysis. Exploration of the AR protein interactome identified G3BP1, SLIRP, and IGFBP5 as AR protein partners which are associated with stress, adaptive metabolic response and estrogen receptor repression.

**Conclusions:** The findings of this study highlight the prognostic potential of cytoplasmic AR immunoreactivity in specific breast cancer subtypes and uncover novel extra-nuclear AR protein interactions that may mediate metabolic adaptations during the development of endocrine-resistance.

## TRANSLATIONAL RELEVANCE

### Is cytoplasmic androgen receptor the ‘tell’ for an estrogen receptor alpha bluff?

Detection of ER protein within the primary tumour plays a major role in dictating breast cancer patient therapy. AR is often detectable in ER positive breast cancers. We have previously reported that elevated AR levels associate with poor outcome to aromatase inhibitor therapy.

In this study, we demonstrate that the localisation of the AR may have a bearing on the activity levels of ER and on clinical outcome. We show cytoplasmic AR to be associated with shorter survival in the Luminal B cohort (p=0.008), an effect only seen in the post-menopausal cohort.

Our study refines the role of AR and specifically its intracellular localisation as a clinically-relevant diagnostic and prognostic marker. Furthermore, our study improves our understanding of the biology of the AR in breast cancer and its role in metabolic adaptations in the context of an androgenic steroid milieu, and identifies new therapeutic targets associated with cytoplasmic AR localisation.

## Introduction

Androgens are involved in the regulation of many cellular processes such as development, transcription and metabolism (reviewed in Bleach & McIlroy (*1*)). Much of our understanding around androgen action is focused upon the ligand driver role of its cognate transcription factor, the androgen receptor (AR). AR actions can be divided into two main pathways, commonly referred to as canonical AR signalling, which involves AR binding to DNA, and non-canonical AR signalling which is not directly dependent on direct AR DNA binding (*2*). In particular, the physiological role of non-genomic AR signalling, particularly in breast cancer, has not been fully elucidated. AR is expressed in up to 99% of hormone receptor positive breast cancer tumours (*3*). AR protein expression, as determined by immunohistochemistry (IHC), is currently the primary criteria for inclusion into breast cancer clinical trials of AR targeting therapies (*4*). However, there are inconsistencies around the cut-point applied, and in addition, various AR antibodies are routinely used to detect expression. Furthermore, clinical trials are exploring the use of both agonists and antagonists of AR with a lack of conclusive studies to indicate which is more likely to offer therapeutic benefit to patient subtypes. It is reported that response to AR-targeted therapies in unselected breast cancer patients is relatively low (*5*). This highlights the lack of clarity and understanding as to the role AR plays in breast cancer across the spectrum of this disease.

Sex steroid production is controlled by a complex network of enzymes present in specific tissues. Of particular importance in breast cancer is the enzyme aromatase which converts androgens to estrogens. Targeted endocrine treatment of postmenopausal breast cancer involves the use of therapies known as aromatase inhibitors (AI) which prevent systemic estrogen synthesis. Studies suggest that due to the mechanism of action of AI therapy, the levels of estrogen detected in breast tumours is almost undetectable (*6*). We hypothesise that adaptive changes due to long-term exposure of breast cancers to the resulting super-abundance of androgens, enables AR to promote growth and survival in the AI resistant setting (*7, 8*). The physiological role of non-genomic AR signalling, particularly in breast cancer, has not been fully elucidated. Recent studies show that non-canonical AR activity is not effectively blocked by AR antagonism, hence it is important to understand the role of the AR outside of its classical transcription factor functions in endocrine resistant breast cancer (*9, 10*). Unlike the genome, the proteome is both dynamic over time and location specific. Steroid nuclear receptors are a prime example of proteins whose function is directly impacted by interactions with other proteins. Studies have uncovered over 350 proteins that interact with ligand activated androgen receptor (AR) (*11*), with a large amount of these proteins involved in gene transcription modulation (*12*). Many studies have focused on understanding the AR interactome in prostate cancer (reviewed (*13*)) but very few studies have explored the AR interactome in breast cancer (*14*).

In this study we analysed the role of AR in AI resistant breast cancer through identifying its interactome. In contrast to other studies which artificially manipulate protein expression (*15*) to obtain a global view of prototypical AR-binding proteins in endocrine resistance, we utilised an AI resistant and sensitive breast cancer cell model with endogenously elevated levels of AR protein (both nuclear and cytoplasmic) to best recapitulate its biological functions in this setting (*16*). The findings of this study highlight novel extra-nuclear AR protein interactions that could aid our understanding of the role played by androgens in metabolic health and the development of endocrine-resistance.

## Results

### Cytoplasmic localisation and impact on survival across breast cancer subtypes

Non-genomic actions of the AR have been well described as potential mediators of resistance in prostate cancer in particular. More recently, Keeming et al (*9*) have also reported non-canonical AR activity as a driver of endocrine resistance in breast cancer. We decided to explore this is in a breast cancer tissue microarray (TMA) previously immunohistochemically (IHC)-stained for AR (*8*). Whilst previous IHC staining of AR intensity focused on nuclear expression, this time, we selectively identified tissue cores with detectible cytoplasmic levels of AR and correlated these with patient outcome. Levels of detectable cytoplasmic AR was determined using Halo digital pathology software (Indica Labs) (Figure 1A).

**Figure 1.**
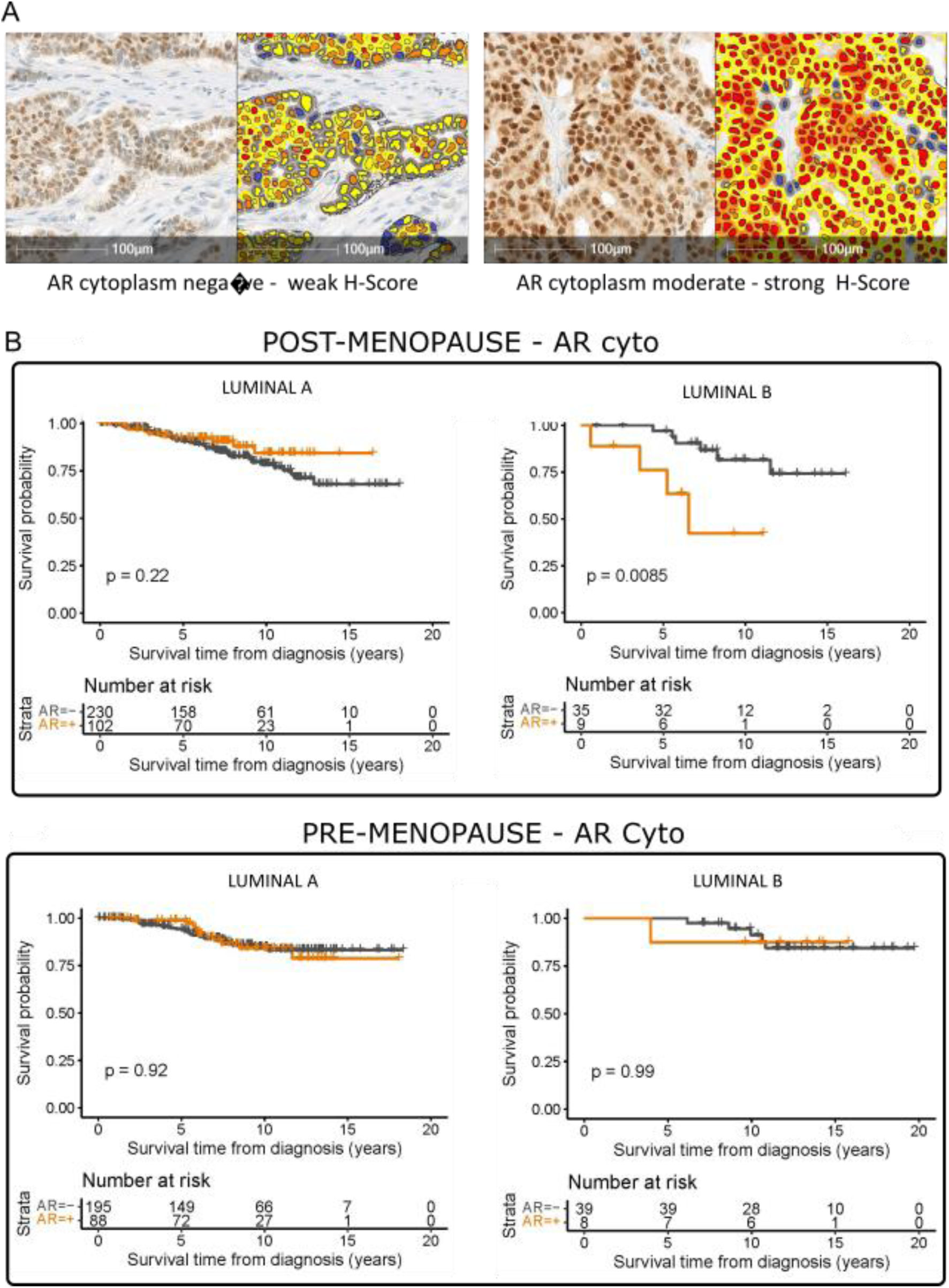
Extra-nuclear detection of cytoplasmic AR associates with poor survival in post-menopausal luminal B breast cancers but has no impact in the pre-menopausal cohort. **A** Representative IHC images of AR staining and the AR Nuc and Cyto mark-up of the same images using HALO digital pathology showing weak-negative and moderate-high staining. Kaplan-Meier plots assessing survival based on detectable AR Cyto staining in breast cancer patients from the CTI-09/07 trial. Top panel – depicting survival in Luminal A and Luminal B subtype stratified by post-menopausal status. Lower panel - depicting survival in Luminal A and Luminal B subtype stratified by pre-menopausal status.

High levels of cytoplasmic AR (>H-score Q3) were significantly associated with poor survival in post-menopausal Luminal B patients (p=0.0085, n=44) but were non-significant when evaluated in this cohort when premenopausal (p=0.99, n=47) (Figure 1 B). Of note, cytoplasmic AR had no impact on luminal A patient cohorts, irrespective of menopausal status. Significantly, cytoplasmic AR was also associated with loss of PR protein expression post-menopause (p=0.008, n=219) (Table 1).

**Table 1.** Clinical and pathologic parameters and their association were evaluated by ^¶^ Fisher exact test [ER^+^, PR^+^, HER2+,recurrence, metastasis, metastatic spread]. and ^Ψ^ Chi-squared test [histology, clinical subtype, histological grade].

|  | Total | Pre-menopausal | % pre-menopausal | Post-menopausal | % post-menopausal | p-value |
| --- | --- | --- | --- | --- | --- | --- |
| Receptor status: AR high | n = 219 | n = 98 | 45 | n = 121 | 55 |  |
| Receptor status: Estrogen <sup>¶</sup> | 219 |  |  |  |  |  |
| ER positive | 207 | 96 | 46 | 111 | 54 | 0.0699 |
| ER negative | 12 | 2 | 16 | 10 | 84 |  |
| Receptor status: Progesterone <sup>¶</sup> | 219 |  |  |  |  |  |
| PR positive | 172 | 85 | 49 | 87 | 51 | 0.0083 |
| PR negative | 47 | 13 | 28 | 34 | 72 |  |
| Receptor status: HER2 <sup>¶</sup> | 219 |  |  |  |  |  |
| HER2 positive | 21 | 10 | 47 | 11 | 53 | 0.820 |
| HER2 negative | 198 | 88 | 44 | 110 | 56 |  |
| Clinical subtype <sup>ψ</sup> | 219 |  |  |  |  | 0.047 |
| Luminal A | 190 | 88 | 46 | 102 | 54 |  |
| Luminal B | 17 | 8 | 47 | 9 | 53 |  |
| HER2 (ER-) | 4 | 2 | 50 | 2 | 50 |  |
| TNBC | 8 | 0 | 0 | 8 | 100 |  |
| Histology <sup>ψ</sup> | 219 |  |  |  |  | 0.192 |
| IDC | 178 | 84 | 47 | 94 | 53 |  |
| ILC | 24 | 9 | 38 | 15 | 62 |  |
| IDC+ILC | 9 | 1 | 11 | 8 | 89 |  |
| Histological Grade <sup>ψ</sup> | 219 |  |  |  |  | 0.740 |
| Grade I | 46 | 22 | 48 | 24 | 52 |  |
| Grade II | 115 | 52 | 45 | 63 | 55 |  |
| Grade III | 56 | 24 | 43 | 32 | 57 |  |
| Recurrence <sup>¶</sup> | 219 | 98 |  | 121 |  |  |
| Yes | 40 | 23 | 58 | 17 | 42 |  |
| No | 179 | 75 | 42 | 104 | 58 | 0.080 |
| Metastasis <sup>¶</sup> | 219 |  |  |  |  |  |
| Yes | 47 | 27 | 57 | 20 | 43 |  |
| No | 172 | 71 | 41 | 101 | 59 | 0.068 |
| Metastatic spread <sup>¶</sup> | 47 | 27 |  | 20 |  |  |
| Local | 7 | 1 | 14 | 6 | 86 |  |
| Distant | 40 | 26 | 65 | 14 | 35 | 0.031 |

#### Androstenedione exposure impacts AR protein levels, cellular localisation and signalling

The cellular location of AR is dynamic depending on cell type, ligand present and biological processes occurring in the cell (*17*). Androstenedione (A4) is the most abundant direct precursor of more potent sex steroids in the postmenopausal setting, and a previous publication from our lab showed a significant increase of A4 levels in AI resistant patients (*8*). As shown in Fig 2A, there was a significant increase in nuclear AR expression in response to A4 treatment in AI resistant MCF7aro-LetR (LetR) cells, however, nuclear ER expression was unchanged (Figure 2A). With this in mind, we devised live-cell time-lapse imaging experiments to establish the cellular location of AR following A4 exposure in LetR cells. To verify that A4 exposure is capable of inducing nuclear translocation of the AR, an additional experiment was conducted to confirm this in the context of GFP-tagged, exogenously over-expressed AR (Figure 2B). It should be noted that the translocation to the nucleus occurs more slowly with A4 (> 1hr) than if cells are exposed to the more potent AR ligand, R1881 (< 10 minutes). Furthermore, using our cell line models of AI resistance, we showed that p-ERK second messenger signalling is increased in a rapid (3-20 minute) manner when cells were exposed to A4, in contrast there was no impact on p-AKT signalling (Figure 2C). Whilst all experiments are conducted in the presence of the aromatase inhibitor letrozole, the cells do express a comprehensive panel of steroidogenic enzymes at a transcript level (See Supplemental Figure 1). Collectively, these data show that A4 and/or its metabolites are capable of triggering abundant expression of AR protein coupled with increased cytoplasmic retention time and that this is likely mediating non-genomic AR signalling.

**Figure 2.**
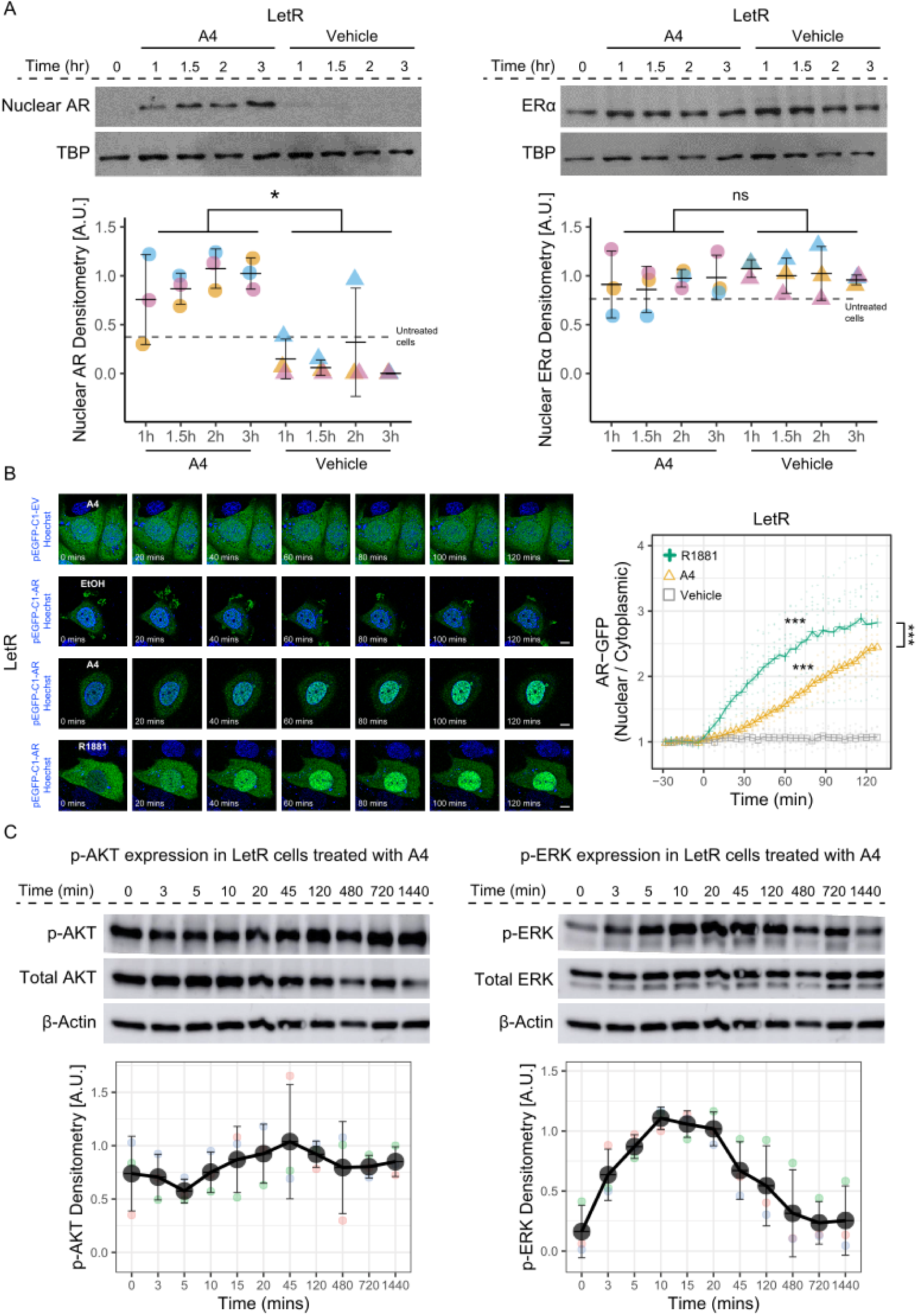
Androstenedione exposure impacts AR protein levels, cellular localisation and signalling. **A** Western blot analysis of nuclear protein lysate was utilised to investigate AR and ER responsiveness to A4 and/or its metabolites in AI resistant LetR cells. Images shows AR and ER protein detected in LetR cells exposed to A4 10^−7^ M or vehicle over a time course of 3 hours. TATA binding protein (TBP) was used as a nuclear lysate loading control. Densitometry analysis of n=3 western blots show AR and ER protein post A4 treatment compared to vehicle (EtOH) normalised to control. Dotted line shows AR nuclear protein expression levels in untreated cells. Densitometry measurements represented as arbitrary units (A.U.). **Two-**way ANOVA analysis was carried out for statistical comparison *P<0.05, non-significant (ns). **B** Representative time-lapse images of LetR cells transfected with pEGFP-C1-AR or empty vector (pEGFP-C1-EV) control plasmid. Both plasmids contain a green fluorescent protein (GFP) tag and the cell nucleus is stained with Hoechst in blue. Top 2 Panels - empty vector control plasmid and LetR transfected with pEGFP-C1-AR plasmid treated with A4 10^−7^ M and EtOH, respectively. Lower 2 panels - Representative time lapse images of LetR cells expressing pEGFP-C1-AR plasmid treated with A4 10-7 M for 3 hours. Representative time lapse images of LetR cells expressing pEGFP-C1-AR plasmid treated with R1881 12.5×10-9 M for 3 hours. These experiments were repeated 3 times with a minimum of 25 cells imaged in each biological replicate. Changes in nuclear/ cytoplasmic ratio over time with A4 and R1881 stimulus (8 cells per group were analysed and ANOVA followed by Tukey’s HSD was performed to assess statistical comparisons, * * * P< 0.001). **C** Western blot analysis was used to evaluate if A4 treatment activates cytoplasmic second messenger signalling proteins in LetR cells over a 24 hour timecourse. Representative image of western blots obtained showing pERK, total ERK, pAKT, total AKT and β-actin control in LetR cells. Quantification of levels of pAKT and pERK protein using densitometry analysis of western blot results at various time points following A4 treatment. Graphed results have been normalised to β--actin loading control. Both graphs are representative of three experimental replicates.

#### Phenotypic responses to A4 observed in AI resistant but not AI sensitive cells

Based on the literature evidence that AR may influence stem cell populations (Barton et al., 2017) (de Kruijff et al., 2019) we postulated that A4 exposure may have a functional impact on the stem cell-like properties of the AI resistant cells. First generation mammosphere formation assesses the ability of the cells to form non-adherent mammosphere structures. These mammospheres can then be dissociated to assess self-renewal capability of the cells, which is indicative of stem cell activity (*18*). Morphological differences in mammosphere formation were observed between parental MCF7 and AI resistant cells. LetR cells formed larger, looser mammosphere structures than isogenic MCF7 cells (Figure 3A). LetR cells in the presence of A4 also formed more mammospheres than MCF7 cells in A4. LetR cells in A4 exhibited increased self-renewal capability compared to their vehicle control and MCF7 cells in A4 (Figure 3B, upper panels). As previous studies from our group have suggested that AI resistant cells may utilise AR as a mechanism of resistance, we decided to explore the possibility that the anti-AR therapy, enzalutamide, may be effective at decreasing stemness in these cells. We have previously shown that enzalutamide decreases LetR but not MCF7 cell viability (*7*). Similarly, enzalutamide treatment decreased the mammosphere formation efficiency of LetR but not MCF7 cells. This result was also mirrored in assessment of self-renewal ability. LetR cells self-renewal competency was decreased significantly. Whilst MCF7 self-renewal also displayed a decrease in the presence of enzalutamide, it was not statistically significant ((Figure 3B, lower panels).

**Figure 3.**
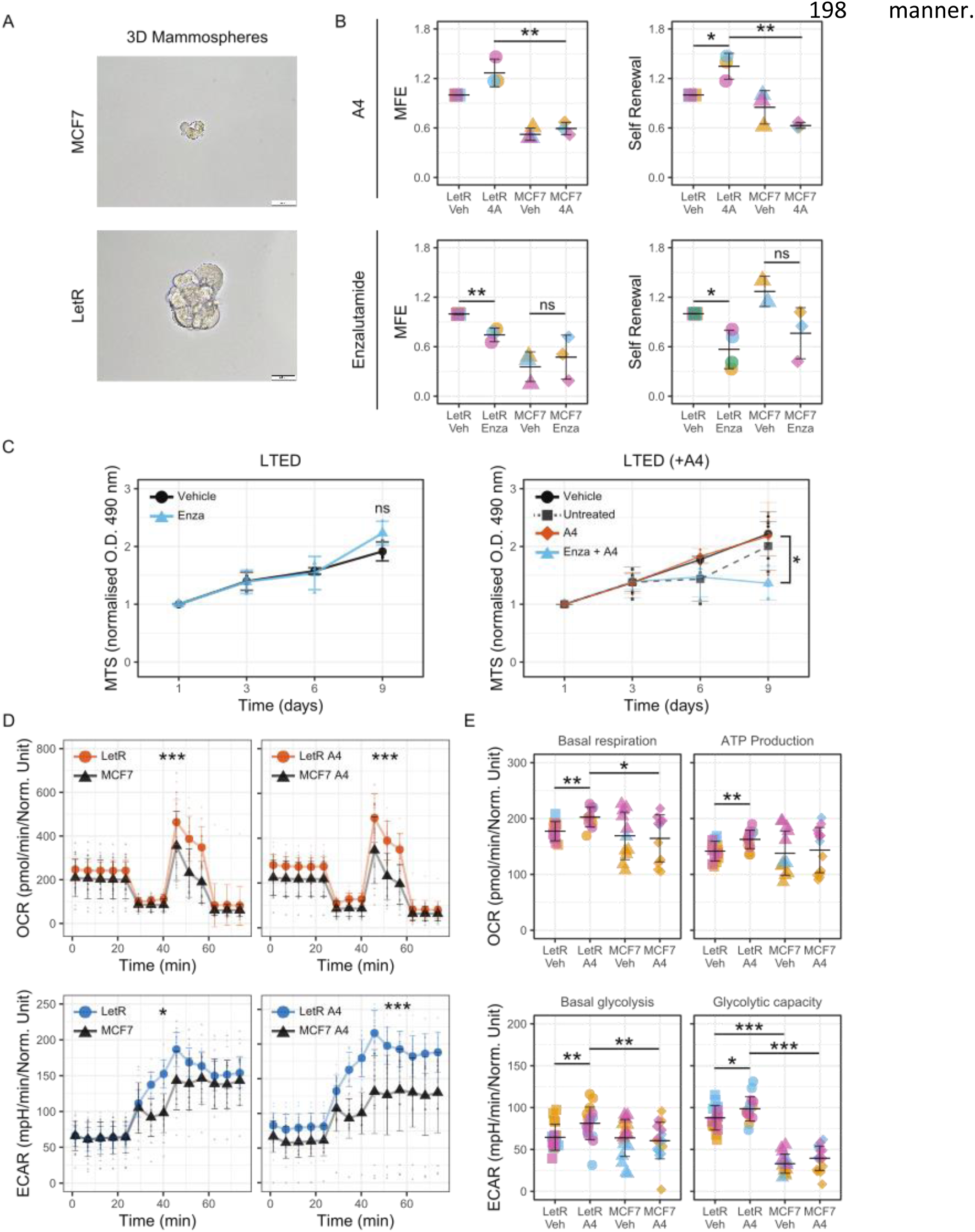
Alterations in the androgenic steroid environment impact cellular response to anti-AR therapy. **A** Representative images of mammosphere structures in LetR and MCF7 cells cultured in ultra-low adherence plates for 6 days. Scale bar = 50 µm. **B** Upper panels - Graphs show mammosphere forming efficiencies (MFE) and percentage self-renewal of LetR and MCF7 cells treated with 10-7 M A4 or vehicle for 6 days. Lower panels - The ability of the anti-AR therapy enzalutamide (10 μM) to inhibit mammosphere formation was assessed in AI resistant LetR cells and parental MCF7 cells after 6 days. The effect of enzalutamide on self-renewal capacity over 6 days in culture was also investigated. Graphs are representative of three experimental replicates. **C** Left hand graph depicts MTS results of LTED cells in media containing no steroids and treated with enzalutamide 10 µM over 9 days. Right hand graph shows MTS assay results from LTED cells grown in A4 10-7 M and simultaneously treated with 10 μM enzalutamide. All graphs are normalised to day 1 reading to account for seeding density. Graphs are representative of n=3 biological replicates. Student’s paired, 2-tailed t-test was carried out on day 6 and day 9 values. **D** Seahorse analysis of mitochondrial stress test in AI resistant LetR and AI sensitive MCF7 cells. Upper panel - Oxygen consumption rate (OCR), a measure of oxphos was evaluated. Results show OCR readings from MCF7 and LetR cells at a basal level and when treated with A4 10-7 M for 24 hours. Lower panel - Extracellular acidification rate (ECAR), a measure of glycolysis was evaluated in LetR and MCF7 cells. Results displayed show basal glycolysis in AI resistant LetR cells versus AI sensitive MCF7 cells and ECAR readings of the cells following 10-7 M A4 treatment for 24 hours. **E** Upper panels - Basal respiration rates in MCF7 and LetR cell lines under A4 or control treatments. Lower panels - Basal glycolysis rates of LetR cell treated with 10-7 M A4 or vehicle control, and MCF7 cells treated with 10-7 M A4 or vehicle control. Graph shows glycolytic capacity of LetR or MCF7 cells under treatment conditions indicated. Time course data was analysed using two-way ANOVA to assess all changes over time and treatments. Bar chart data is presented as StDev two-tailed unpaired t-test, *P<0.05., ** p<0.01, non-significant (ns).

We then wanted to further explore the impact that the steroid environment has on anti-AR treatment response by testing the impact of the steroid environment on long term estrogen deprived (LTED) cells. LTED cells cultured in the absence of steroids are unresponsive to enzalutamide treatment. However, when LTED cells are cultured over the course of the experiment in growth medium supplemented with A4, a significant decrease in viability in response to enzalutamide is evident (Figure 3C). Of note, as there was no increase in LTED viability in the presence of A4, and enzalutamide only decreases viability in the presence of A4, it suggests A4 may enhance cell survival in a non-mitogenic manner.

#### Androgen excess increases the metabolic fitness and plasticity of breast cancer cells

We have previously observed alteration in cell size and morphology between AI resistant and sensitive cells when exposed to a hyperandrogen environment (*7*). Given the strong links between AR and metabolism (review by (*19*)) we first looked at mitochondrial mass and potential using the isogenic MCF7 and LetR cell line models. A4 exposed LetR exhibited donut-shaped mitochondrial morphology, increased mitochondrial mass, and increased mitochondrial potential (Supplemental Figure 2A and B). We then assessed Oxygen Consumption Rate (OCR) and ExtraCellular Acidification Rate (ECAR) as a measure of mitochondrial respiration and glycolysis in AI resistant and sensitive cells (Figure 3D). After subtracting the levels of non-mitochondrial respiration in all samples, AI resistant LetR cells treated with A4 were found to have increased levels of basal mitochondrial respiration compared to vehicle control and MCF7 cells. LetR cells exposed to A4 also had the highest levels of adenosine triphosphate (ATP) synthesis compared to vehicle treated LetR or MCF7 (Figure 3E, upper panels) suggesting that A4 treatment is driving processes that require energy production in the cells. Interestingly A4 did not cause any significant changes in AI sensitive MCF7 cells providing evidence for altered response to the steroid environment. Maximal respiration capacity was higher in LetR A4 treated cells over MCF7 A4 treatment. In addition, the level of spare respiratory capacity was almost doubled in LetR cells compared to MCF7 suggesting LetR cells are much more capable of responding to increased energy demand or conditions of stress (Supplemental Figure 2C).

The shift in cellular respiration to glycolysis when challenged with specific inhibitors of mitochondrial respiration was also assessed. Results indicate that AI resistant LetR cells have enhanced glycolysis under basal conditions compared to MCF7 cells. Furthermore, when cells are treated with oligomycin which inhibits mitochondrial respiration, a much greater adaptability to shift to glycolysis was noted in LetR cells. MCF7 cells displayed an immediate increase in glycolysis however were unable to sustain the response. In contrast LetR cells continually increased glycolysis over time. LetR A4 treated cells also displayed higher <u>glycolytic capacity and</u> maximum glycolytic rate compared to MCF7 A4 and vehicle treated, although this was only slightly increased over LetR vehicle treated control cells (Figure 3E, lower panels, and Supplemental Figure 2C).

#### AR interactome in AI resistant breast cancer

We then used a proteomic based approach to investigate the impact of A4 and/or its metabolites on AR protein interactions to try and elucidate the non-mitogenic impact of A4 on cell viability. AR is known to interact with an extensive array of proteins in a variety of cell types and treatment conditions in order to exert its pleiotropic effects. In this experiment we set out to investigate the unique AR interactome that supervenes in AI treatment resistance. As breast cancer is primarily assessed in the clinic through immunohistochemical staining, novel protein biomarkers could provide clinically applicable means by which to assess AI therapy treatment response at point of diagnosis. In order to address the question of what proteins interact uniquely with AR in the AI resistant environment we utilised our *in vitro* cell models (MCF7 and MCF7Aro-LetR) and carried out an AR co-immunoprecipitation (Co-IP) mass spectrometry experiment. Cell lines were treated with A4 10^−7^ M, cholesterol (chol) 10^−7^ M or ethanol (EtOH) as lipid and vehicle control respectively, for 3 hours. Letrozole 10^−6^ M was added in the presence of A4 to prevent its conversion to estrone and also in the control treated samples. Cholesterol is the precursor from which all steroids are derived and therefore, was included as an additional lipid control. Co-IP experiments often result in non-specific proteins binding to the matrix. Here, we pre-cleared the lysate by incubation with control protein A/G beads to remove proteins with an affinity for the column matrix (*20*). After the cells were lysed and AR was immunoprecipitated, the samples were sent for electrospray ionisation (ESI) MS/MS analysis - Experimental workflow was summarized Figure 4A. Analysis of results involved eliminating proteins that were common to interacting with AR in all samples. The end goal was to identify AR protein complexes only present in LetR cells under A4 exposure.

**Figure 4.**
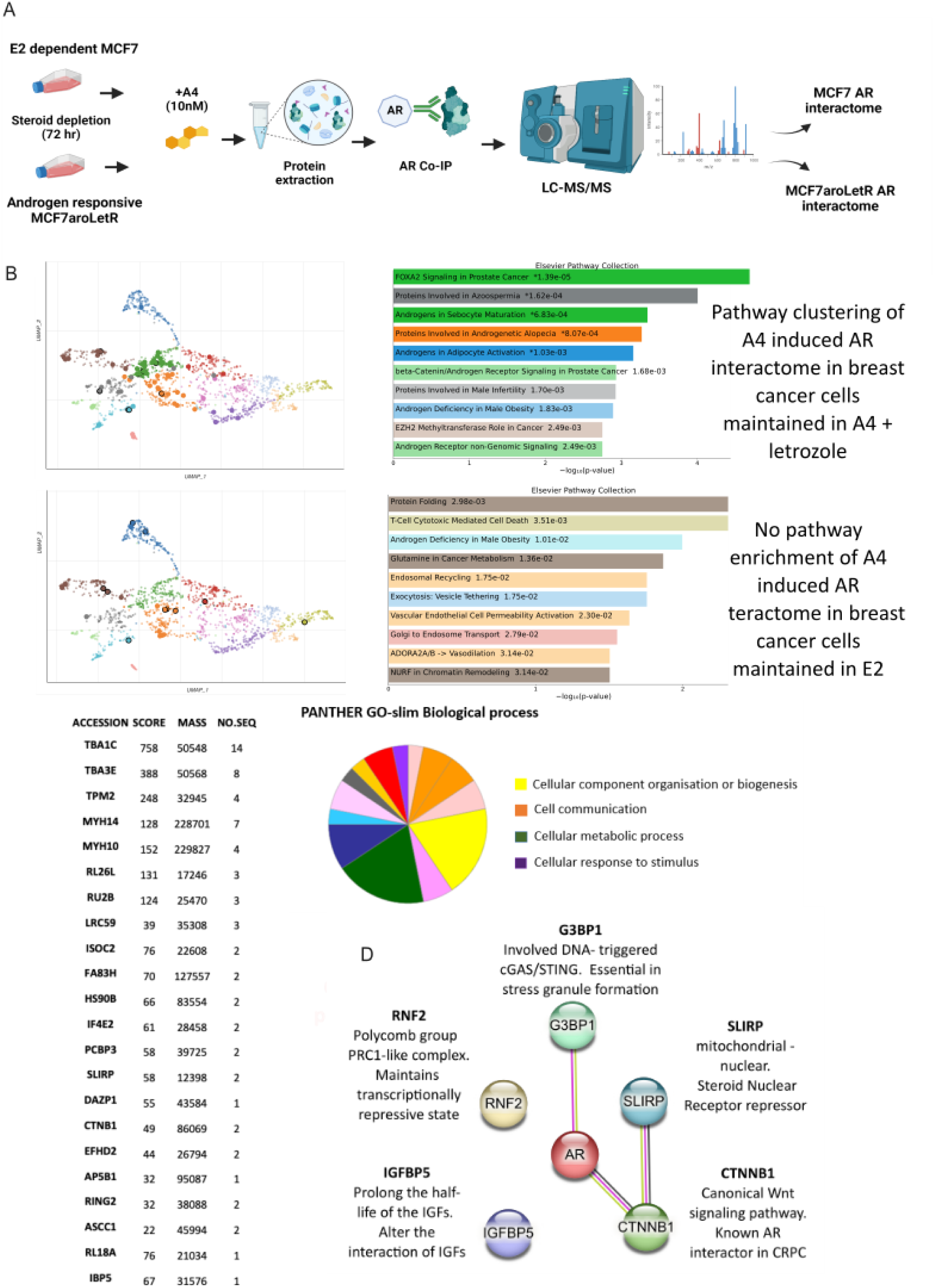
AR interactome in AI resistant breast cancer. **A** Schematic of the mass-spectrometry workflow used to explore the AR interactome. MCF7 (maintained in estrogen) and MCF7Aro-LetR (maintained in androgen) were treated with A4 vs. vehicle control for 3 hours. Following co-IP, samples were isolated prior to electrospray Ionisation (ESI) MS/MS analysis. **B** A global analysis of pathway enrichment was carried out to identify networks associated with the AR interactome. In cells maintained under longterm androgen and letrozole there was significant enrichment of key pathways post A4 exposure, primarily associated with non-genomic AR signalling (significant pathways in bold coloured bars with white font, non-significant pathways are in pastel with black font). There were no pathways significantly enriched by short-term exposure to A4 in the cells maintained under estrogen. **C** AR interacting proteins unique to LetR cells (A4 vs. EtOH) were further shortlisted based on number of peptides (≥2) or Mascot score (≥67). GO biological processes of AR interacting proteins unique to LetR cells were uploaded into Cytoscape (Bingo plugin) to determine statistically overrepresented biological processes in this group of proteins (pie-chart and listed). **D** STRING network analysis was carried out on the top interacting proteins determined to be strong candidate proteins identified as differential AR interactors with A4 treatment in the AI resistance setting.

783 unique AR interacting proteins were identified under all treatment conditions in LetR cells. 1073 proteins were identified in MCF7 cells under all treatments. Of these proteins 255 proteins were exclusive to the AI resistant LetR cells and 545 proteins were identified as exclusive to MCF7 cells. Proteins numbers interacting with AR identified in the AI resistant LetR cells included: 83 proteins under cholesterol treatment, 64 proteins in vehicle and 59 proteins with A4 treatment. The number of proteins identified as interacting with AR in MCF7 cells included: 83 under cholesterol treatment, 208 under vehicle conditions and 140 with A4 treatment (See Supplemental Table 1 & 2 for A4 interacting proteins in MCF7 and LetR respectively).

A number of known AR interacting protein were identified compared with AR RIME (Rapid Immunoprecipitation of Endogenous proteins) and AR interacting proteins and co-regulators (Supplemental Table 3). The number of AR co-factors identified was relatively low compared to previously published AR RIME data (*21*). However, it should be noted AR RIME has only been conducted in prostate cancer cells and therefore many differential cofactors may be present compared to breast cancer cells. Also as the experimental aim was to identify both cytoplasmic and nuclear protein interacting AR proteins no enrichment of cellular fractions was conducted. The sheer abundance of proteins in whole cell lysate may have masked the detection of lesser abundant cofactor proteins interacting with AR.

Pathway analysis (https://maayanlab.cloud/Enrichr/) showed distinct differences in the clustering of the AR protein partners between the MCF7 and LetR interactome. The LetR AR interactome exhibited a preponderance of pathways associated with genomic and non-genomic androgen signalling including FOXA2 signalling in prostate cancer, androgenic maturation of sebocytes and beta catenin/ AR signalling in prostate cancer. In contrast, within the MCF7 AR interactome subset there was indistinct clustering with no significant pathway enrichment (Figure 4B). Cytoscape was then used to decipher the potential functions of proteins specifically interacting with AR under A4 stimulus in the AI resistant LetR cells. Swissprot identifiers are shown alongside their Mascot score and respective numbers of unique peptides detected. The resulting PANTHER gene ontology analysis showed that a number of cell processes were associated including: cell communications, metabolic processes, component organisation, organelle biogenesis and response to stimulus.

Additional validity tests were carried out using an online platform: contaminant repository for affinity purification (CRAPome) (https://www.crapome.org) to ensure bona fide AR protein interactions. This resource combines negative controls from multiple sources of affinity purification mass spectrometry experiments can greatly improve the chances of eliminating background contaminants (*22*). Using the number of experiments in which a protein was detected, combined with the average spectral scores, the likelihood of contamination was determined for each protein (Supplemental Figure 3B). In total seven unique A4 associated AR protein interactions in LetR cells were selected as candidates for downstream validation, IGFBP5, SLIRP, CTNBN1 and RNF2 were identified as unlikely to be contaminants. CTNNB1 ((Catenin beta-1) adherent junctions, cell adhesion, WNT signalling pathway) is the only known AR protein partner. All other protein are novel putative interactors: IGFBP5 (Insulin-like growth factor-binding protein 5) – Involved in the regulation of IGF signalling. SLIRP (SRA stem-loop-interacting RNA-binding protein, mitochondrial) – Coactivator/repressor of nuclear receptors. RNF2 (E3 ubiquitin-protein ligase RING 2) – chromatin binding and ubiquitin protein transferase activity (Figure 4D).

### Validatory experiments confirm that AR interacts with SLIRP and IGFBP5 which co-localise in proximity to the nuclear membrane and within nuclear foci under A4 treatment in AI resistant cells

Based on the top proteins identified as differential A4 driven AR interactors in LetR cells, and coupled with knowledge of the literature on known AR interacting proteins both SLIRP and IGFBP5 were validated by co-IP followed by western blot analysis (Figure 5A). SLIRP has previously been identified as an AR interacting protein in prostate cancer (*23*). IGFBP5 is a novel AR interactor identified in this study but it has previously been identified as playing a role in breast cancer (*24*).

**Figure 5.**
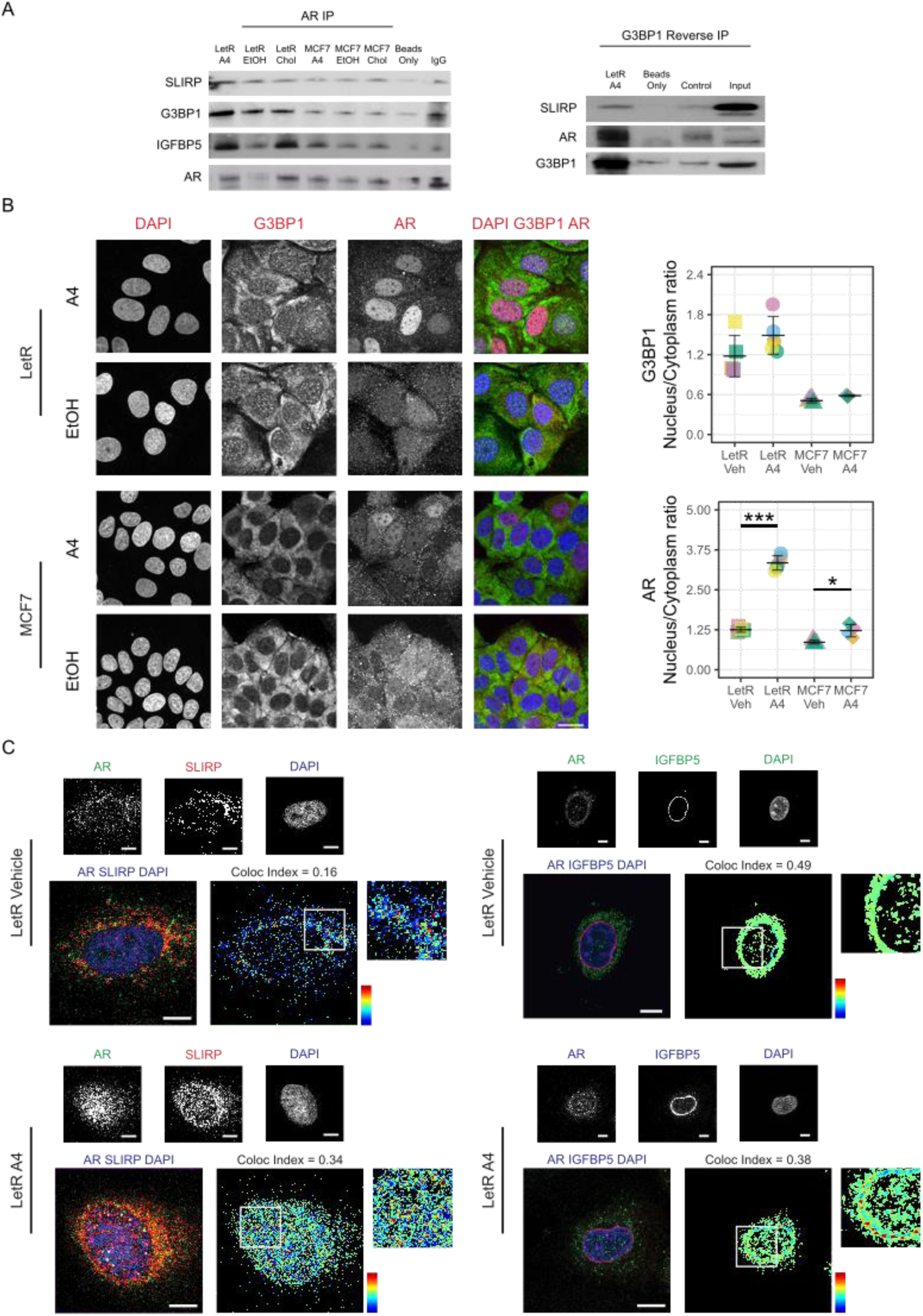
Validation of AR protein interactions in AI resistant and AI sensitive cells. **A** Western blotting was used to confirm AR immunoprecipitation followed by confirmation of the presence of interacting proteins SLIRP, G3BP1 and IGFBP5. 1000 µg of protein was used as the co-IP input for LetR and MCF7 exposed to A4, EtOH, cholesterol. The first 6 wells contain AR immunoprecipitate and interacting proteins with equalised quantities of input protein and antibody. The final two wells are negative controls with beads only and corresponding IgG control. B Representative confocal microscope imaging of LetR cells and MCF7 cells for AR and G3BP1 protein localization analysis with A4 treatment or vehicle treatment. Graphs show nuclear/cytoplasm ratio of AR protein and G3BP1 protein expression analysed by mean fluorescence intensity of AR/G3BP1 fluorescence in nucleus /cytoplasm. **C** LetR cells were treated with EtOH (upper panels) or A4 (lower panels) for 3 hours before being fixed and stained. Mid plane of 3D confocal microscopy imaging depict AR and SLIRP expression detected by AR, SLIRP/IGFBP5 and DAPI nuclear stain. Images show overlay of AR antibody staining (green) with SLIRP/IGFBP5 antibody staining (red). Colocalization Colormap ImageJ plugin was used to assess colocalisation and calculate colocalisation idex, colour bar indicates high (red) and low (blue) colocalisation. Scale bar is 10 μm.

For validation of the AR protein interactome data we also decided to take forward a high scoring protein defined as having 10 unique peptides across all treatment conditions but with higher detection as an AR interactor in AI resistant LetR cells compared to MCF7 cells. From the MS results, G3BP1 displayed a quantitative value normalised to total spectra which was almost 2-fold higher in the LetR cells under all treatment conditions compared to the MCF7 (Supplemental Figure 3B). Reverse IP of G3BP1 in the LetR cells under the same experimental conditions as outlined above confirmed interaction with both AR and SLIRP (Figure 5A). Further immunofluorescent imaging analysis showed increased nuclear/ cytoplasmic localisation of AR and G3BP1 in LetR cells when exposed to A4 versus control conditions. Low levels of nuclear/ cytoplasmic G3BP1 was notable in the MCF7 cells under all experimental conditions (Figure 5B).

To further confirm the interactions between AR with SLIRP and IGFBP5, 3D confocal microscopy was utilised. For consistency with the co-IP mass spectrometry experimental set up, cells were treated for 3 hours with control vehicle (Figure 5C – upper panels) or A4 (Figure 5C - lower panels) before fixation and staining. Antibody detected endogenous AR showed minimal AR in the nucleus under vehicle treatment. SLIRP expression was detected in the cytoplasm of the cells and a small amount in the nucleus under vehicle stimulation (Figure 5C, left panels). We also observed a weak colocalisation between SLIRP and AR in vehicle treated cells. In contrast, under A4 treatment conditions the majority of AR was detected in the peri-nuclear/ nuclear region of LetR cells. Similarly, an increase in SLIRP expression in the nucleus of LetR cells was also evident, and there was twice as much colocalisation between AR and SLIRP as compared to vehicle. These results indicate that AR and SLIRP co-localise both inside the nucleus and in the cytoplasm of LetR cells under A4 treatment. IGFBP5 protein demonstrated peri-nuclear localization post A4 treatment. Comparison of the images under each treatment condition, combined with our previous evidence demonstrating AR nuclear translocation upon A4 stimulation indicates that AR and IGFBP5 interact in a peri-nuclear location in AI resistant LetR cells when A4 is present.

## Discussion

This study demonstrates distinct roles of the AR in the setting of Luminal A tumours and the more insidious Luminal B subtype but only post-menopause. With increasing age, the levels of the prohormones produced in our bodies steadily decline with the exception of some adrenal-specific androgens whose levels remain elevated and may have relevance for the development of diseases of metabolism including cancers (*25, 26*). In previous studies we have reported an association between higher AR signalling coupled with elevated adrenal androgens and poor response to endocrine therapy in post-menopausal breast cancers (*8*). Understanding how a preponderance of these adrenal androgens influence AR activity and interactions could therefore provide insight as to how they impact breast cancer development and therapy resistance. It is widely accepted that disorders of androgen excess in women can result in lifelong alterations in metabolism and are associated with the development of type 2 diabetes, metabolic dysfunction associated fatty liver disease (MAFLD) and polycystic ovary syndrome (PCOS) (*27–29*). Furthermore, extra-nuclear AR has been reported to modulate mitochondrial activity and to form complexes, responsible for the activation of transmembrane adenylate cyclase (*30, 31*). Breast cancer risk strongly associates with metabolic syndromes, which poses the question of a causative role for androgen excess in the development of this disease and subsequent therapy resistance.

The concept that systemic androgen excess may trigger non-genomic actions of the AR has been highlighted in previous research (*32*). Herein, we have confirmed elevation of AR protein and activation of second messenger signalling cascades as a result of adrenal androgen excess and we have also shown this to be associated with poor outcome in a breast cancer patient cohort. Moreover, we observed changes in receptor kinetics with AR cytoplasmic retention time increased with exposure to less potent androgenic steroids that are known to dominate in metabolic diseases of female androgen excess (*27, 33*). The significance of androgen excess is further emphasized when we show that absence of estrogen from the system is not sufficient to replicate the post-menopausal state, illustrated by data showing the differential impact of AR blockade in the absence or presence of androgens. We also show the potent impact of A4 (and/or its derivatives) on both breast cancer cell respiration and glycolysis when cells are chronically exposed to A4. This data can help address some of the deficits in our understanding of the dimorphic effects of androgens in males and females (*34–36*). Whilst potent androgens have been reported to boost mitochondrial health in men and in rodent models (*37*); studies in women with diseases of androgen excess, such as PCOS, demonstrate malformed mitochondrial structures. In our models we observed increased mitochondrial mass in cells exposed to androgen, with large numbers of swollen and fragmented mitochondria. From these data we surmise that the effects of androgens on mitochondrial function in women is dependent, not simply upon the potency of the ligand, but also their relative abundance. Our model recreates the distortion of mitochondrial morphology observed in other diseases of androgen excess in females but the energy profile is more similar to that observed in the condition of obesity/ nutrient overload (*36*). The link between androgens, lipid trafficking and mitochondrial bioenergetics is long established and could in part explain the highly energetic state of the breast cancer cells exposed to A4, and also must be considered in the context of the mammary adipose-rich environment. This is therefore the first study to look at the impacts of excess androgens with high bioavailability in relation to mitochondrial activity and which neatly recapitulates the clinical scenario.

Given that we observe different phenotypes in AI resistant and sensitive cells induced by the altered steroid environment (*7*), and coupled with the large amount of data on the role of AR in metabolism we postulated that metabolic alterations may be induced in AI resistant models. In support of this theory AI resistant cells that are maintained under androgen excess were found to have increased metabolic fitness characterised by both increased mitochondrial and glycolytic capacity compared to sensitive cells. Breast cancer cells are known to display metabolic plasticity (*38*) and this is exemplified in our AI resistant LetR cells. Furthermore, metabolic plasticity is believed to be a mechanism by which cancer cells avoid programmed cell death and enable tumour progression (*39*). Our results demonstrate a greater ability in LetR cells to switch to glycolysis than MCF7 cells. Moreover, endocrine treatment resistance can alter metabolic processes, as demonstrated by tamoxifen resistant cells that display gene patterns associated with mitochondrial dysfunction and increased mitochondrial activity (*40*). In light of our previously published study showing reliance on AR in AI resistant cells and the recently established roles of AR in mediating cellular metabolism (*19, 30*), we hypothesised that the metabolic phenotype observed in AI resistant cells may involve AR and interacting protein partners.

Using a mass-spectrometry based approach we identified a number of high confidence interactors (G3BP1, SLIRP and IGFBP5) in addition to beta-catenin, a known AR protein partner that interacts with AR in castrate-resistant prostate cancer (*41*). Of note, all putative interactors are reported to be primarily localised to extra-nuclear sites within the cell such as mitochondria (SLIRP), cytoplasmic vesicles (IGFBP5) and cytosol (G3BP1). Our data show enhanced AR protein interaction within both cytoplasmic and nuclear compartments when cells are exposed to androstenedione. This is a significant finding as both SLIRP and G3BP1 have been shown to act as nuclear receptor repressors and activators of mRNA decay respectively (*42, 43*), moreover, the polycomb repressor complex 1 component, RING2 was also identified as a high confidence AR partner. This is in line with other studies that have shown AR to antagonise ER genomic activity and also our own translational findings, wherein we reported high levels of AR and serum androgens to be associated with decreased ESR1 gene activity (*8, 44*). Indeed the heightened levels of G3BP1:AR interaction in the AI resistant cells suggests deployment of pro-survival stress response mechanisms linked to selective partitioning of transcription and associated metabolic rewiring (*45*). That these putative adaptive AR protein interactions only occur as a result of longterm androgen exposure highlights the significance of exploring the impact of hormones as systemic, long-lasting signals within mammalian systems. Another validated AR interactor, IGFBP5, is associated with inflammation and fibrosis and is also reported to both promote and reinforce senescence via auto and paracrine mechanisms (*46*). Whilst, our data have not fully elucidated the mechanism of IGFBP5:AR engagement, there does appear to be a degree of perinuclear localisation. This is in keeping with other studies reporting extranuclear localisation of IGFBP5 to be pro-migratory in contrast to its growth inhibitory nuclear role (*24*). Previously published data from our group has reported morphological and transcriptional alterations in response to adrenal androgens that are strongly indicative of altered metabolism and the expression of a senescence-associated secretory phenotype (SASP) (*47*). Most notably, we identified prosaposin to be associated with poor outcome to endocrine therapy, which is significant as it has long been associated with SASP via re-direction of lipid mediated lysosomal - mitochondria signalling and enhanced metabolism (*48*). Collectively, this data leads us to hypothesise that longterm exposure to androgen excess can distort cancer cell metabolism and facilitate the development of a more aggressive phenotype. It is however important to stress that further exploratory experiments will be required to fully elucidate the mechanisms underlying all these interactions.

Tantalisingly, these data raise the possibility of cytoplasmic AR acting as in indicator of impaired ER function, as such, furthering our understanding of the extra-nuclear interactions of AR will provide invaluable insights as to how best to target Luminal B breast cancers prone to therapy resistance. This patient population is particularly poorly served by current therapeutic regimens and thus utilising digital pathology approaches to map AR localisation could pinpoint vulnerable individuals at a much early stage of their disease. In conclusion, extrapolation of the findings of this study provide much needed insights into the role of AR under conditions of androgen excess in breast cancer and potentially underlies the development of a range of chronic disorders of metabolism in women.

## Materials and Methods

### Indica Halo digital pathology tissue microarray analysis

We accessed a TNBC data from a previous study in which mouse anti-human monoclonal AR primary antibody (AR-318-L-CE, Leica Biosystems) was used to detect AR in a tissue microarray (TMA) of primary breast carcinomas (Bleach et al., 2021). Digital whole slide images (WSI) of the TMAs were created by scanning the slides at X40 magnification using a high power digital scanner (Glissando, Objective Imaging). The HALO AI (Indica Labs, Albuquerque, NM, USA) DenseNet v2 classifier was trained using manual annotation by the Pathologist (KS) to segment tumour epithelium from tumour stroma. The tissue classification was followed by AR quantification of DAB (diaminobenzidine) IHC staining using the HALO IHC multiplex and TMA modules, assessing quantification of AR receptor staining in the cytoplasmic cellular compartment of tumour epithelium only. All digital markup results were confirmed by the Pathologist (KS).

### Transfections using Electroporation

LetR and MCF7 cells were transfected with a pEGFP-C1-AR or pEGFP-C1 empty vector plasmid. Nucleofection (Lonza nucleofector 2b) was used to transfect the plasmid into the cells following protocol from Amaxa® Cell Line Nucleofector® Kit V. This is a technique which uses electrical pulses to create small pores in the cell membrane in order to enhance plasmid entry into the cells. Cells were harvested by trypsinization and counted using countess automated cell counter (Invitrogen). 1 × 10^6^ cells were centrifuged at 90 x g for 10 minutes and then resuspended in 100 μl nucleofector solution V (Lonza). The cells were combined with 2000 ng plasmid DNA and placed in a specially designed cuvette. The appropriate programme (P-20) was selected on the nucleofector device and the cuvette inserted. When complete the sample was transferred to a 35 × 10 mm glass bottom WillCo-dish® (WillCo Wells B.V, Netherlands) pre-incubated at 37°C. After 24 hours transfection, efficiency was established by fluorescent microscopy observing cells expressing green fluorescent protein.

### Live cell imaging

Cells were incubated in 3% PRFMEM medium containing 1 µg/ml Hoechst 33258 (Sigma Aldrich, Darmstadt, Germany) for three hours. After medium was replaced, WillCo-dishes containing AR-GFP-transfected and Hoechst-stained cells were mounted on LSM 710 Confocal laser-scanning microscope (Carl Zeiss Ltd, Cambridge, UK) equipped with a 63×/1.4 NA Plan-Apochromat oil immersion objective and a microscope incubator chamber (set to 37 °C with 5% CO_2_) (Pecon, Erbach, Germany). Hoechst and GFP were excited using 405 nm (1 % laser power, detection range 415-494 nm) and 488 nm (5 % laser power, 490-544 nm) lasers, respectively. Differential interference contrast (DIC), Hoechst and GFP images were recorded at a resolution of 1024 × 1024 pixels for four hours at four minute intervals. A4 10^−7^ M or EtOH 0.01% treatment was added after ∼30 minutes of background measurement. Image processing (background subtraction and median filtering) was carried out using ImageJ (version 1.52i, Wayne Rasband, NIH, Bethesda, MD, USA).

#### Mammosphere formation assay

First generation mammospheres are used to determine mammosphere forming efficiency which is an indication of stem cell activity in cell lines and experiemnst were set up follwong the protocol optimised by Shaw et al., 2012. Briefly, on day 6 the number of mammospheres over 50 µm diameter was counted using CellSens Olympus software. Analysis of mammosphere forming efficiency (MFE) was calculated by MFE = (No. of mammospheres per well / No. of cells seeded per well) × 100. These mammospheres were then passaged to establish self-renewal abilities. Self-renewal = Total number of 2° mammospheres formed / Total number of 1° mammosphere formed. Carried out as per published protocol (*18*).

#### Co-Immunoprecipitation (Co-IP)

MCF7Aro-LetR and MCF7 cells were seeded at 1.5 ×10^6^ and 3.3 ×10^6^ cells respectively into cell culture dish 150 x 20 mm (Sarstedt). They were steroid depleted for 72 hrs in 3% PRFMEM media, with media replenished every day to remove steroids. At 80% confluence cells were treated with A4 10^−7^ M, cholesterol 10^−7^ M or vehicle (ethanol) 0.01% for 3 hours. The plates were then placed on ice, washed and scraped into ice-cold PBS with added protease inhibitor (Merck, NJ). Protein extraction was carried out using the Thermo Fisher Scientific (MA, USA) NE-PER Nuclear and Cytoplasmic extraction kit and quantified using BCA assay (Thermo Fisher Scientific, MA, USA) as outlined above. 1000 µg of whole cell lysate was used for the IP. AR protein was immunoprecipitated using the Pierce Crosslink Immunoprecipitation kit (Thermo Fisher Scientific, MA, USA) according to manufacturer’s instructions. All steps were carried out at 4 °C except were stated. A column with irreversibly bound AR antibody (441): sc-7305 (10 μg) or Mouse IgG Sc-2025 (10 μg) to A/G agarose beads using disuccinimidyl suberate (DSS) was made. Pre-cleared protein lysate was incubated with antibody-crosslinked resin in the column with end-over-end mixing overnight at 4 °C. This included an IgG column and a bead only column as negative control for non-antibody specific protein binding complexes. Protein was eluted from columns using a low pH elution buffer and analysed by western blotting.

### Gel digest and ESI MS/MS analysis

Mass spectrometry acquisition was performed at the proteomics facility at St. Andrew’s University, Scotland. The eluted protein was run approximately 1 cm on a precast Bolt Bis-Tris Plus Gels (Thermo Fisher Scientific). The gel band was excised and cut into pieces (∼1-2 mm^3^) and placed in low protein binding tubes. The samples were then subjected to in-gel digestion, using a ProGest Investigator in-gel digestion robot (Genomic Solutions) using standard protocols (*49*). Briefly, the gel pieces were destained by repeated washing with acetonitrile, and 20 mM Ammonium Bicarbonate, and subjected to reduction (20 mM DTT) and alkylation (50 mM Iodoacetamide) before digestion with trypsin at 37°C (ratio ∼50:1 protein:enzyme). Volumes used were sufficient to cover the gel pieces in each case. The peptides were extracted with 5% formic acid and concentrated down to 20 µl using a SpeedVac (ThermoSavant).

A portion of the resultant peptides were then injected on an Acclaim PepMap 100 C18 trap and an Acclaim PepMap RSLC C18 column (Thermo Fisher Scientific), using a nanoLC Ultra 2D plus loading pump and nanoLC as-2 autosampler (Eksigent). The peptides were held on the trap and washed for 20 mins to waste before switching in line with the column and were eluted with a gradient of increasing acetonitrile, containing 0.1 % formic acid (2-20% acetonitrile in 90 min, 20-40% in a further 30 min, followed by 98% acetonitrile to clean the column, before re-equilibration to 2% acetonitrile). The eluate was sprayed into a TripleTOF 5600+ electrospray tandem mass spectrometer (ABSciex, Foster City, CA) and analysed in Information Dependent Acquisition (IDA) mode, performing 120 msec of MS followed by 80 msec MSMS analyses on the 20 most intense peaks seen by MS. The MS/MS data file generated via the ‘Create mgf file’ script in PeakView (Sciex) was analysed using the Mascot search algorithm (Matrix Science), against the SwissProt database (May2018) restricting the search to Homo sapiens (20,350 sequences). An addition search against NCBI database (Aug2016) all species (93,482,448 sequences) was carried out to check for contaminants. Settings include: trypsin as the cleavage enzyme and carbamidomethylation of cysteine, methionine oxidation as a variable modifications. The peptide mass tolerance was set to 20 ppm and the MS/MS mass tolerance to ± 0.05 Da.

### Network and Gene Ontology analysis

Files containing proteins identified through MASCOT software, with a false discovery rate (FDR) of ≤ 5% were calculated using a randomised database approach, and SwissProt human only searches were analysed. Results were shortlisted using RStudio. This involved removing proteins common to each subgroup. Pathway enrichment analysis was performed on the AR interactome unique to the LetR cells (54 proteins) and MCF7 (131 proteins) respectively (See supplemental Tables 1 & 2) using Enrichr (https://maayanlab.cloud/Enrichr/) (*50*). Cytoscape was then used to decipher the potential functions of proteins interacting with AR in the AI resistant LetR cells compared to the AI sensitive MCF7 cells under all treatment conditions. To gain an understanding of the biological processes that differ in AI sensitive cells compared to AI resistant cells Cytoscape with the Bingo plug-in was again utilised.

### Stratification of proteins for downstream validation

In order to reduce the number of potential false identifications for downstream validation by western blotting, we set the minimum requirement of a Mascot score (Ion score) of ≥ 67 (See Supplemental Figure 3 for Ion score generation graph) or having at least 2 significant peptide matches. Mascot/Ion scores are generated by an algorithm in the programme which assesses number of total, unique and significant peptides. Using this stringent selection criteria, 17 proteins remained in the cholesterol treated group, 13 in the vehicle and 7 unique to A4 treatment as interacting with AR in LetR cells. The same cut-off was applied to the MCF7 cells and the results showed 12 proteins unique to interacting with AR under cholesterol treatment, 35 under vehicle treatment and 27 with A4 treatment. These proteins represent the unique AR interactome in AI resistant and sensitive cells under each treatment condition.

### Immunofluorescent staining

2,500 cells were seeded per chamber in coverglass plates and treated with A4 10^−7^ M or vehicle (0.01% EtOH) for 3 hours. The cells were fixed in 100% methanol for 10 minutes and washed with PBS. 0.1% triton-x-100 (Sigma Aldrich) was used to permeabilise the cells. Cells were blocked with 10% (v/v) goat serum in 5% BSA (w/v) (Sigma Aldrich) for one hour. Following a wash step, AR antibody (sc-816 - rabbit) dilution 1:50 and IGFBP5 primary antibody (sc-515116 - mouse) dilution 1:150 or SLIRP antibody (ab51523) dilution 1:200 was added in 10% human serum (v/v) BSA/PBS (w/v) for 90 minutes. Conjugated secondary antibody against each primary antibody host species was added at 1:200 dilution. This was incubated in darkness at room temperature for one hour. Finally, a blue fluorescent probe called DAPI which binds to the DNA was placed on the sample for 30 seconds in order to label the nucleus. One chamber received an IgG control at the same concentration as the antibodies and another chamber acted as a secondary only control to account for optimal antibody concentration and check specificity of staining.

Cells were then placed on LSM 710 Confocal laser-scanning microscope (Carl Zeiss Ltd, Cambridge, UK) fitted with a 63×/1.4 NA Plan-Apochromat oil immersion objective. 405 nm, 488 nm, 561 nm and 647 nm lasers were used to excite DAPI, Alexa Fluor 488, Alexa Fluor 561 and Alexa Fluor 647 conjugated secondary antibodies with detection ranges of 415-494 nm, 490-544 nm, 560-628 nm and 638-728 nm respectively. Z-stacks were taken by recording series of optical slices with 1 µm thickness at 1024 × 1024 pixels throughout the entire cells. Further image processing (background subtraction, median filtering and construction of orthogonal views) was carried out using ImageJ (version 1.52i, Wayne Rasband, NIH, Bethesda, MD, USA).

### TMRE Flow Cytometry

The mitochondrial membrane potential (MMP or Δψm) was determined using Tetramethylrhodamine (TMRE), a fluorescent dye that accumulates within active mitochondria. 1 × 10^5^ cells were harvested and incubated with 100 nM TMRE for 30 min at 37 °C and 5% CO_2_. Cells were then washed with ice-cold PBS and pelleted at 1200 rpm for three minutes at 4 °C. Cell pellets were then re-suspended in 1 µg/ml PBS-DAPI solution and filtered through a 40 µm filer to ensure cells were non-aggregated and in a single cell suspension. The fluorescent intensity of TMRE was detected using the Atunne NxT Flow Cytometer (ThermoFisher Scientific), and graphs/ illustration were acquired using the FlowJo software (BD Bioscience). Data were obtained from 5 independent experiments, and the mean was calculated relative to the control using Graphpad Prism.

### Seahorse Mitostress Assay

A Seahorse XF Analyzer (Agilent technologies, CA, USA) was used to measure cellular metabolism in live cells through detection of oxygen consumption rate (OCR) and extracellular acidification rate (ECAR). OCR and ECAR reading are used to assess levels of mitochondrial respiration and glycolysis occurring under pre-determined conditions. In this experiment Seahorse XF Cell Mito Stress Test kit protocol was followed. MCF7, LetR, ZR75.1 and ZRR cells were serum starved for 24 hours for synchronisation prior to seeding into seahorse experimental plate (Agilent technologies). 15,000 MCF7, 7,500 LetR, 25,000 ZR75.1 and 25,000 ZRR cells were seeded into their respective wells in 3% PRFMEM media (Sigma Aldrich). No cells were seeded into the outer wells of the plate, these were filled with media to help prevent evaporation. Four corner wells which contained media only were used as reference wells. After allowing cells to adhere for 24 hours they were treated with 10^−7^M 4AD or .01% ethanol vehicle control with letrozole 10^−6^ M present in both conditions. Treatments were carried out in 6 technical replicates. An XE96 four port sensor seahorse cartridge was hydrated with 200 μl calibration buffer (Agilent technologies) per well overnight before running the experiment on the Seahorse machine. Seahorse XF base medium, unbuffered without phenol red was used with the addition of 5.6 mM glucose (Sigma) and 2 mM glutamine (Sigma). This concentration of glucose and L. glutamine was chosen as it is the level in 3% PRFMEM media used as baseline media for all experiments in this study. The pH of this seahorse assay media was adjusted to 7.4. Stimulants and inhibitors of mitochondrial and glycolysis pathways are injected through ports into each well. The following drugs were provided as part of the Seahorse XF Cell Mito Stress Test kit (Agilent technologies) and diluted to give 1 μM Oligomycin, 0.5 μM Carbonyl cyanide 4-(trifluoromethoxy)phenylhydrazone (FCCP) or 0.5 μM rotenone/antimycin as a final concentration after injection into each well from equilibrated port sensor cartridge. To perform the assay the culture medium on the cells was replaced with 150 μl seahorse assay medium and the plate was placed in a CO_2_ free incubator at 37 °C for 1 hour. The calibration plate was placed into the XF96 analyzer seahorse machine approximately 15 minutes prior to the addition of the Seahorse experiment plate. When calibration was completed the lower plate of the sensor cartridge was removed and the seahorse plate containing the cells was inserted. On completion of the assay the seahorse plate containing the cells was removed and kept to quantify cells in each well.

### Statistical analysis

Kaplan-Meier estimates for breast cancer-specific overall survival (OS) and progression-free survival (PFS) were computed based on AR-cyto protein expression in a TMA cohort consisting of n = 875 cases. The OS time for each patient was determined in years from the date of diagnosis (or, when the diagnosis date was unavailable, the primary surgery date) until the date of death or censoring. Progression-free survival (PFS) was calculated in years from the date of diagnosis (or primary surgery) to the occurrence of the first recurrence surgery, date of death, or censoring. Statistics of DAB H-score distribution across each TMA cohort were obtained using R base summary and histogram functions. Cut-off points to estimate survival were determined by selection of the upper quartile (75 % percentile, DAB H-score = 41.69). Kaplan-Meier survival estimates were generated using the survfit and the ggsurvplot functions in the R packages survival (v3.4-0) and survminer (v0.4.9). Statistical significance between the means of two groups was tested using unpaired two-tailed t test. ANOVA with Tukey’s post-hoc test was used for comparing three or more groups. R and GraphPad Prism software were used to perform statistical comparison tests and preparation of graphs. Figures were prepared using Adobe Illustrator or Inkscape. Asterisks represent significance with *, P < 0.05; **, P < 0.01; and ***, P < 0.001. All experiments were repeated at least three times unless otherwise indicated.

## Ethics and consent

Written and informed consent was acquired prior to collection of patient tumor tissue under The Royal College of Surgeons Institutional Review Board–approved protocol (#13/09:CTI 09/07).

## Acknowledgments

M. Mcllroy is supported by Beaumont Hospital Cancer Research and Development Trust, project number: 2077.

