## Supplemental Files for "Androgen receptor interactions provide insight into steroid mediated metabolic shifts in endocrine resistant breast cancer"

**Supplemental Figures**

Supplemental Figure 1

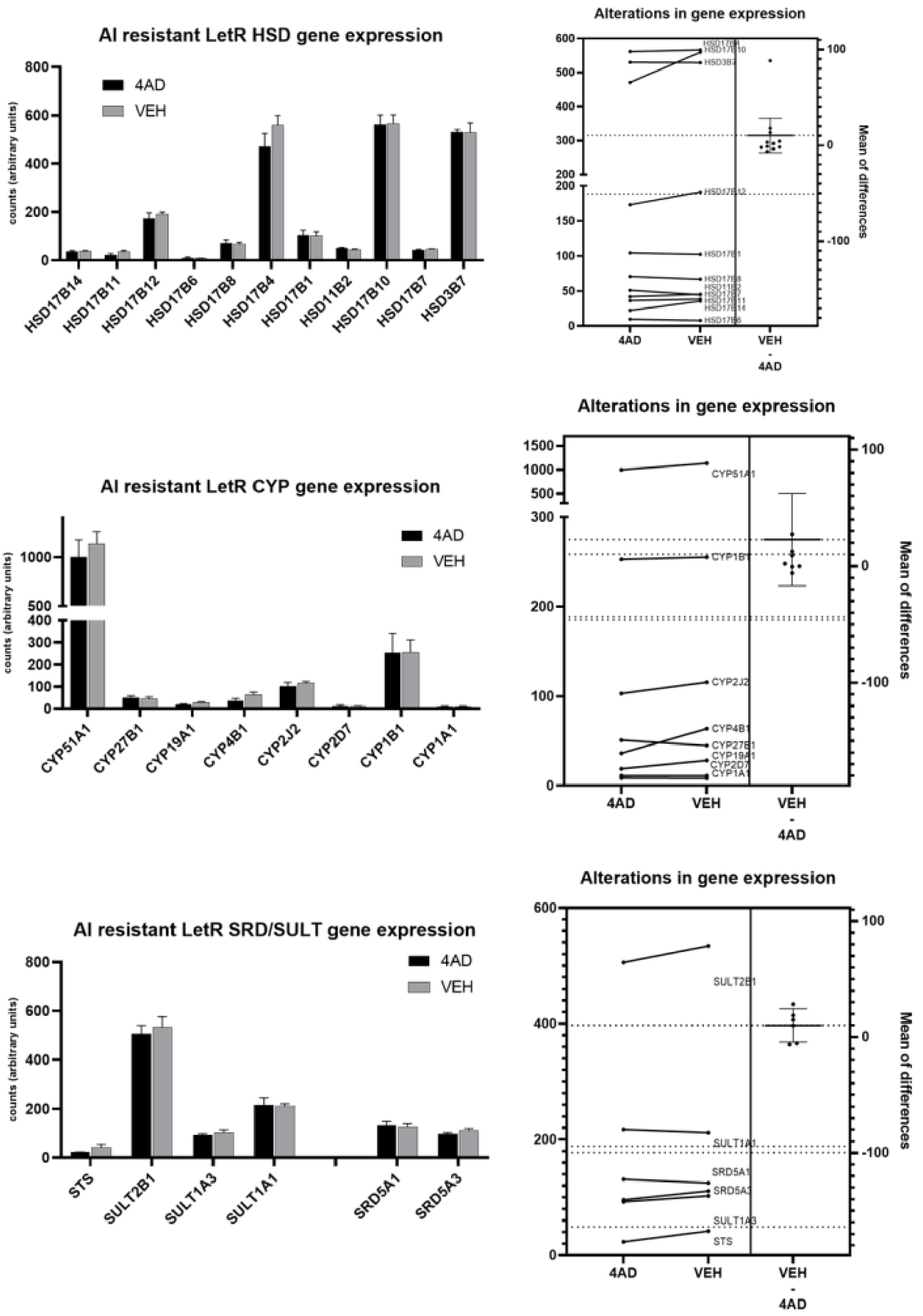

Supplemental Figure 2

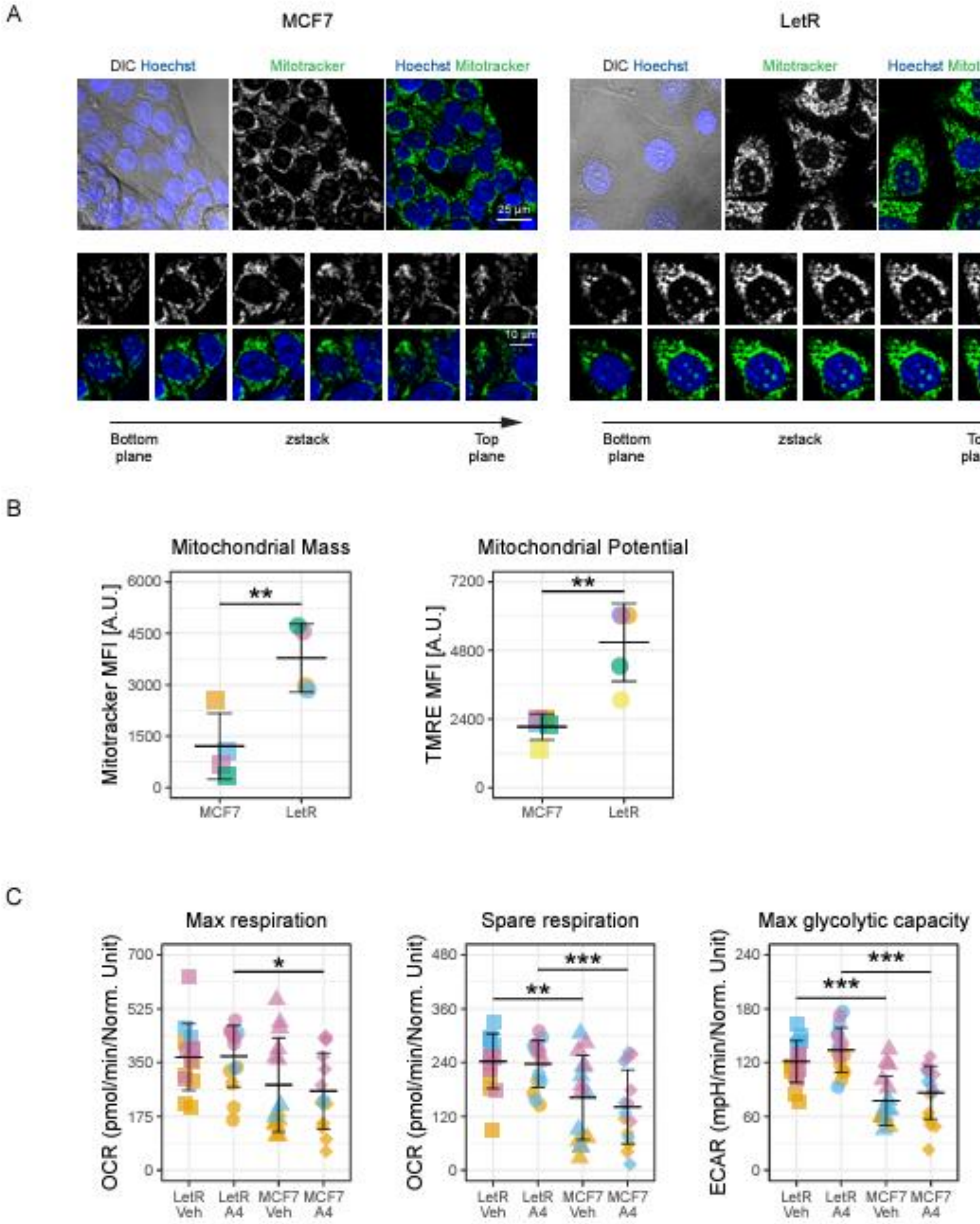

Supplemental Figure 3

| Protein | Score | Mass | Matches | Sequences | Coverage |
|---|---|---|---|---|---|
| SLIRP | 58 | 12398 | 2 | 2 | 22% |
| IGFBP5 | 69 | 31576 | 1 | 1 | 4% |
| G3BP1 | 598 | 52189 | 20 | 12 | **35%** |

```
  1 [illegible]
 51 [illegible]
101 [illegible]
```

```
  1 MVLLTAVLLL LAAYAGPAQS LGSFVHCEPC DEKALSMCPP SPLGCELVKE
 51 PGCGCCMTCA LAEGQSCGVY TERCAQGLRC LPRQDEEKPL HALLHGRGVC
101 LNEKSYREQV KIERDSREHE EPTTSEMAEE TYSPKIFRPK HTRISELKAE
151 AVKKDRRKKL TQSKFVGGAE NTAHPRIISA PEMRQESEQG PCRRHMEASL
201 QELKASPRMV PRAVYLPNCD RKGFYKRKQC KPSRGRKRGI CWCVDKYGMK
251 LPGMEYVDGD FQCHTFDSSN VE
```

```
  1 MVMEKPSPLL VGREFVRQYY TLLNQAPDML HRFYGKNSSY VHGGLDSNGK
 51 PADAVYGQKE IHRKVMSQNF TNCHTKIRHV DAHATLNDGV VVQVMGLLSN
101 NNQALRRFMQ TFVLAPEGSV ANKFYVHNDI FRYQDEVFGG FVTEPQEESE
151 EEVEEPEERQ QTPEVVPDDS GTFYDQAVVS NDMEEHLEEP VAEPEPDPEP
201 EPEQEPVSEI QEEKPEPVLE ETAPEDAQKS SSPAPADIAQ TVQEDLRTFS
251 WASVTSKNLP PSGAVPVTGI PPHVVKVPAS QPRPESKPES QIPPQRPQRD
301 QRVREQRINI PPQRGPRPIR EAGEQGDIEP RRMVRHPDSH QLFIGNLPHE
351 VDKSELKDFF QSYGNVVELR INSGGKLPNF GFVVFDDSEP VQKVLSNRPI
401 MFRGEVRLNV EEKKTRAARE GDRRDNRLRG PGGPRGGLGG GMRGPPRGGM
451 VQKPGFGVGR GLAPRQ
```

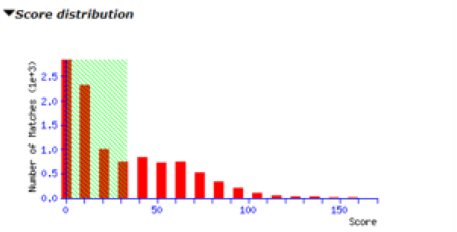

**Peptide score distribution.** Ions score is −10 log(P), where P is the probability that the observed match is a random event.
There are **3,771** peptide matches above identity threshold and **4,778** matches above homology threshold for **30,150** queries. On average, individual ions scores **> 33** (beyond green shading) indicate **identity or extensive homology** (p<0.05).

| Accession | Description | Likely to be contamination |
|---|---|---|
| TBA3E/TUBA3E | Tubulin alpha-3E chain | High |
| TPM2 | Tropomyosin beta chain | Medium |
| IGFBP5 | Insulin-like growth factor-binding protein 5 | Low |
| SLIRP | SRA stem-loop-interacting RNA-binding protein, mitochondrial | Low |
| CTNB1/CTNNB1 | Catenin beta-1 | Low |
| RING2/RNF2 | E3 ubiquitin-protein ligase RING 2 | Low |
| AP5B1 | AP-5 complex subunit beta-1 | Unknown |

**Supplemental Table 1 MCF7 only MS data – proteins with ≥2 peptides or MASCOT score ≥67**

| Accession | Score | Mass | No. of sig. matches | No. of sig sequences | emPAI | Description |
|---|---|---|---|---|---|---|
| POTEE | 3885 | 122882 | 78 | 10 | 0.53 | POTE ankyrin domain family member E OX=9606 GN=POTEE PE=2 SV=3 |
| KRT86 | 1639 | 55120 | 63 | 26 | 8.24 | Keratin, type II cuticular Hb6 OX=9606 GN=KRT86 PE=1 SV=1 |
| POTEI | 1825 | 122858 | 42 | 8 | 0.36 | POTE ankyrin domain family member I OX=9606 GN=POTEI PE=3 SV=1 |
| K1H2 | 709 | 51793 | 36 | 7 | 0.89 | Keratin, type I cuticular Ha2 OX=9606 GN=KRT32 PE=2 SV=3 |
| KRT81 | 294 | 56832 | 36 | 3 | 0.28 | Keratin, type II cuticular Hb1 OX=9606 GN=KRT81 PE=1 SV=3 |
| K2C78 | 238 | 57629 | 34 | 2 | 0.18 | Keratin, type II cytoskeletal 78 OX=9606 GN=KRT78 PE=1 SV=2 |
| BAZ1A | 568 | 180246 | 32 | 2 | 0.05 | Bromodomain adjacent to zinc finger domain protein 1A OX=9606 GN=BAZ1A PE=1 SV=2 |
| KRT37 | 538 | 51084 | 32 | 4 | 0.45 | Keratin, type I cuticular Ha7 OX=9606 GN=KRT37 PE=1 SV=3 |
| KRT34 | 969 | 50818 | 24 | 17 | 3.84 | Keratin, type I cuticular Ha4 OX=9606 GN=KRT34 PE=1 SV=2 |
| PERI | 568 | 53732 | 22 | 3 | 0.3 | Peripherin OX=9606 GN=PRPH PE=1 SV=2 |
| PRKDC | 512 | 473749 | 19 | 18 | 0.2 | DNA-dependent protein kinase catalytic subunit OX=9606 GN=PRKDC PE=1 SV=3 |
| OPHN1 | 76 | 92210 | 17 | 1 | 0.05 | Oligophrenin-1 OX=9606 GN=OPHN1 PE=1 SV=1 |
| ACTBM | 559 | 42331 | 11 | 4 | 0.56 | Putative beta-actin-like protein 3 OX=9606 GN=POTEKP PE=5 SV=1 |
| HS71L | 492 | 70730 | 11 | 7 | 0.6 | Heat shock 70 kDa protein 1-like OX=9606 GN=HSPA1L PE=1 SV=2 |
| RL4 | 241 | 47953 | 8 | 6 | 0.8 | 60S ribosomal protein L4 OX=9606 GN=RPL4 PE=1 SV=5 |
| K1C25 | 216 | 49858 | 8 | 3 | 0.46 | Keratin, type I cytoskeletal 25 OX=9606 GN=KRT25 PE=1 SV=1 |
| U17L1 | 54 | 60692 | 8 | 1 | 0.08 | Ubiquitin carboxyl-terminal hydrolase 17-like protein 1 OX=9606 GN=USP17L1 PE=3 SV=1 |
| RS8 | 357 | 24475 | 7 | 6 | 2.16 | 40S ribosomal protein S8 OX=9606 GN=RPS8 PE=1 SV=2 |
| ADT1 | 256 | 33271 | 7 | 6 | 1.33 | ADP/ATP translocase 1 OX=9606 GN=SLC25A4 PE=1 SV=4 |
| MIC60 | 146 | 84026 | 7 | 7 | 0.48 | MICOS complex subunit MIC60 OX=9606 GN=IMMT PE=1 SV=1 |
| F135A | 39 | 171614 | 7 | 1 | 0.03 | Protein FAM135A OX=9606 GN=FAM135A PE=1 SV=2 |
| COQ9 | 37 | 35658 | 7 | 1 | 0.14 | Ubiquinone biosynthesis protein COQ9, mitochondrial OX=9606 GN=COQ9 PE=1 SV=1 |
| HS105 | 218 | 97716 | 6 | 6 | 0.34 | Heat shock protein 105 kDa OX=9606 GN=HSPH1 PE=1 SV=1 |
| UN45A | 172 | 104266 | 6 | 6 | 0.31 | Protein unc-45 homolog A OX=9606 GN=UNC45A PE=1 SV=1 |
| RL3 | 133 | 46365 | 5 | 5 | 0.66 | 60S ribosomal protein L3 OX=9606 GN=RPL3 PE=1 SV=2 |
| SNED1 | 34 | 158206 | 5 | 1 | 0.03 | Sushi, nidogen and EGF-like domain-containing protein 1 OX=9606 GN=SNED1 PE=2 SV=2 |
| ACD11 | 202 | 88026 | 4 | 4 | 0.24 | Acyl-CoA dehydrogenase family member 11 OX=9606 GN=ACAD11 PE=1 SV=2 |
| RCN1 | 180 | 38866 | 4 | 4 | 0.62 | Reticulocalbin-1 OX=9606 GN=RCN1 PE=1 SV=1 |
| H11 | 167 | 21829 | 4 | 3 | 0.9 | Histone H1.1 OX=9606 GN=HIST1H1A PE=1 SV=3 |
| FLNC | 117 | 293407 | 4 | 4 | 0.07 | Filamin-C OX=9606 GN=FLNC PE=1 SV=3 |
| WDR1 | 111 | 66836 | 4 | 3 | 0.24 | WD repeat-containing protein 1 OX=9606 GN=WDR1 PE=1 SV=4 |
| MYH1 | 107 | 223976 | 4 | 2 | 0.04 | Myosin-1 OX=9606 GN=MYH1 PE=1 SV=3 |
| U5S1 | 101 | 110336 | 4 | 4 | 0.19 | 116 kDa U5 small nuclear ribonucleoprotein component OX=9606 GN=EFTUD2 PE=1 SV=1 |
| FA98B | 95 | 37566 | 4 | 4 | 0.65 | Protein FAM98B OX=9606 GN=FAM98B PE=1 SV=1 |
| RAB14 | 95 | 24110 | 4 | 3 | 0.79 | Ras-related protein Rab-14 OX=9606 GN=RAB14 PE=1 SV=4 |
| AP2M1 | 94 | 49965 | 4 | 4 | 0.46 | AP-2 complex subunit mu OX=9606 GN=AP2M1 PE=1 SV=2 |
| NAT10 | 90 | 116569 | 4 | 4 | 0.18 | RNA cytidine acetyltransferase OX=9606 GN=NAT10 PE=1 SV=2 |
| U520 | 88 | 246006 | 4 | 4 | 0.08 | U5 small nuclear ribonucleoprotein 200 kDa helicase OX=9606 GN=SNRNP200 PE=1 SV=2 |
| MATR3 | 87 | 95078 | 4 | 4 | 0.22 | Matrin-3 OX=9606 GN=MATR3 PE=1 SV=2 |
| TPM4 | 73 | 28619 | 4 | 4 | 0.93 | Tropomyosin alpha-4 chain OX=9606 GN=TPM4 PE=1 SV=3 |
| LIPS | 73 | 117323 | 4 | 1 | 0.04 | Hormone-sensitive lipase OX=9606 GN=LIPE PE=1 SV=4 |
| RLA0L | 71 | 34514 | 4 | 2 | 0.31 | 60S acidic ribosomal protein P0-like OX=9606 GN=RPLP0P6 PE=5 SV=1 |
| RS9 | 63 | 22635 | 4 | 4 | 1.29 | 40S ribosomal protein S9 OX=9606 GN=RPS9 PE=1 SV=3 |
| SHE | 37 | 54259 | 4 | 1 | 0.09 | SH2 domain-containing adapter protein E OX=9606 GN=SHE PE=1 SV=1 |
| HS74L | 149 | 95479 | 3 | 3 | 0.16 | Heat shock 70 kDa protein 4L OX=9606 GN=HSPA4L PE=1 SV=3 |
| AP1B1 | 130 | 105482 | 3 | 3 | 0.14 | AP-1 complex subunit beta-1 OX=9606 GN=AP1B1 PE=1 SV=2 |

| | | | | | | |
|---|---|---|---|---|---|---|
| NOP2 | 128 | 89589 | 3 | 2 | 0.11 | Probable 28S rRNA (cytosine(4447)-C(5))-methyltransferase OX=9606 GN=NOP2 PE=1 SV=2 |
| RS10L | 128 | 20279 | 3 | 3 | 0.99 | Putative 40S ribosomal protein S10-like OX=9606 GN=RPS10P5 PE=5 SV=1 |
| NOP56 | 125 | 66408 | 3 | 3 | 0.24 | Nucleolar protein 56 OX=9606 GN=NOP56 PE=1 SV=4 |
| RL10 | 120 | 25044 | 3 | 3 | 0.75 | 60S ribosomal protein L10 OX=9606 GN=RPL10 PE=1 SV=4 |
| RBM10 | 98 | 103811 | 3 | 3 | 0.15 | RNA-binding protein 10 OX=9606 GN=RBM10 PE=1 SV=3 |
| RPN1 | 88 | 68641 | 3 | 3 | 0.23 | Dolichyl-diphosphooligosaccharide--protein glycosyltransferase subunit 1 OX=9606 GN=RPN1 PE=1 SV=1 |
| HNRL1 | 85 | 96250 | 3 | 3 | 0.16 | Heterogeneous nuclear ribonucleoprotein U-like protein 1 OX=9606 GN=HNRNPUL1 PE=1 SV=2 |
| ELAV1 | 76 | 36240 | 3 | 3 | 0.48 | ELAV-like protein 1 OX=9606 GN=ELAVL1 PE=1 SV=2 |
| MYH7B | 68 | 226960 | 3 | 2 | 0.04 | Myosin-7B OX=9606 GN=MYH7B PE=1 SV=4 |
| IQGA1 | 65 | 189761 | 3 | 3 | 0.08 | Ras GTPase-activating-like protein IQGAP1 OX=9606 GN=IQGAP1 PE=1 SV=1 |
| PICAL | 63 | 70881 | 3 | 3 | 0.22 | Phosphatidylinositol-binding clathrin assembly protein OX=9606 GN=PICALM PE=1 SV=2 |
| MY18A | 62 | 234168 | 3 | 3 | 0.06 | Unconventional myosin-XVIIIa OX=9606 GN=MYO18A PE=1 SV=3 |
| 1433E | 58 | 29326 | 3 | 3 | 0.62 | 14-3-3 protein epsilon OX=9606 GN=YWHAE PE=1 SV=1 |
| DSRAD | 58 | 137178 | 3 | 3 | 0.11 | Double-stranded RNA-specific adenosine deaminase OX=9606 GN=ADAR PE=1 SV=4 |
| PRP6 | 57 | 107656 | 3 | 3 | 0.14 | Pre-mRNA-processing factor 6 OX=9606 GN=PRPF6 PE=1 SV=1 |
| KRT82 | 50 | 57985 | 3 | 2 | 0.18 | Keratin, type II cuticular Hb2 OX=9606 GN=KRT82 PE=1 SV=3 |
| ERR1 | 46 | 46108 | 3 | 1 | 0.11 | Steroid hormone receptor ERR1 OX=9606 GN=ESRRA PE=1 SV=3 |
| DDX21 | 44 | 87804 | 3 | 3 | 0.18 | Nucleolar RNA helicase 2 OX=9606 GN=DDX21 PE=1 SV=5 |
| KTN1 | 39 | 156464 | 3 | 3 | 0.09 | Kinectin OX=9606 GN=KTN1 PE=1 SV=1 |
| TAD2B | 36 | 49010 | 3 | 1 | 0.1 | Transcriptional adapter 2-beta OX=9606 GN=TADA2B PE=1 SV=2 |
| SOS2 | 34 | 154251 | 3 | 1 | 0.03 | Son of sevenless homolog 2 OX=9606 GN=SOS2 PE=1 SV=2 |
| PI3R4 | 25 | 154318 | 3 | 1 | 0.03 | Phosphoinositide 3-kinase regulatory subunit 4 OX=9606 GN=PIK3R4 PE=1 SV=3 |
| DKC1 | 152 | 58094 | 2 | 2 | 0.18 | H/ACA ribonucleoprotein complex subunit 4 OX=9606 GN=DKC1 PE=1 SV=3 |
| RBM39 | 125 | 59628 | 2 | 2 | 0.17 | RNA-binding protein 39 OX=9606 GN=RBM39 PE=1 SV=2 |
| EBP2 | 121 | 34887 | 2 | 2 | 0.31 | Probable rRNA-processing protein EBP2 OX=9606 GN=EBNA1BP2 PE=1 SV=2 |
| RAI3 | 120 | 40624 | 2 | 2 | 0.26 | Retinoic acid-induced protein 3 OX=9606 GN=GPRC5A PE=1 SV=2 |
| HORN | 104 | 283140 | 2 | 2 | 0.03 | Hornerin OX=9606 GN=HRNR PE=1 SV=2 |
| DDX54 | 94 | 98819 | 2 | 2 | 0.1 | ATP-dependent RNA helicase DDX54 OX=9606 GN=DDX54 PE=1 SV=2 |
| KR131 | 94 | 19505 | 2 | 2 | 0.61 | Keratin-associated protein 13-1 OX=9606 GN=KRTAP13-1 PE=2 SV=2 |
| ENPL | 92 | 92696 | 2 | 2 | 0.11 | Endoplasmin OX=9606 GN=HSP90B1 PE=1 SV=1 |
| 1433T | 90 | 28032 | 2 | 2 | 0.4 | 14-3-3 protein theta OX=9606 GN=YWHAQ PE=1 SV=1 |
| CPT1A | 90 | 88995 | 2 | 1 | 0.05 | Carnitine O-palmitoyltransferase 1, liver isoform OX=9606 GN=CPT1A PE=1 SV=2 |
| RS26 | 89 | 13292 | 2 | 2 | 1.01 | 40S ribosomal protein S26 OX=9606 GN=RPS26 PE=1 SV=3 |
| PTH2 | 88 | 19466 | 2 | 2 | 0.62 | Peptidyl-tRNA hydrolase 2, mitochondrial OX=9606 GN=PTRH2 PE=1 SV=1 |
| MYL9 | 88 | 19871 | 2 | 1 | 0.27 | Myosin regulatory light polypeptide 9 OX=9606 GN=MYL9 PE=1 SV=4 |
| PK1IP | 85 | 44506 | 2 | 1 | 0.11 | p21-activated protein kinase-interacting protein 1 OX=9606 GN=PAK1IP1 PE=1 SV=2 |
| FXR2 | 85 | 74520 | 2 | 2 | 0.14 | Fragile X mental retardation syndrome-related protein 2 OX=9606 GN=FXR2 PE=1 SV=2 |
| PSIP1 | 85 | 60181 | 2 | 2 | 0.17 | PC4 and SFRS1-interacting protein OX=9606 GN=PSIP1 PE=1 SV=1 |
| MTA2 | 84 | 75717 | 2 | 2 | 0.13 | Metastasis-associated protein MTA2 OX=9606 GN=MTA2 PE=1 SV=1 |
| TBL2 | 83 | 50393 | 2 | 2 | 0.21 | Transducin beta-like protein 2 OX=9606 GN=TBL2 PE=1 SV=1 |
| KRA31 | 83 | 11558 | 2 | 2 | 1.23 | Keratin-associated protein 3-1 OX=9606 GN=KRTAP3-1 PE=1 SV=1 |
| TCPA | 79 | 60819 | 2 | 2 | 0.17 | T-complex protein 1 subunit alpha OX=9606 GN=TCP1 PE=1 SV=1 |
| H31T | 78 | 15613 | 2 | 2 | 0.81 | Histone H3.1t OX=9606 GN=HIST3H3 PE=1 SV=3 |
| THEM6 | 78 | 24135 | 2 | 2 | 0.48 | Protein THEM6 OX=9606 GN=THEM6 PE=1 SV=2 |
| ARP3B | 78 | 48090 | 2 | 2 | 0.22 | Actin-related protein 3B OX=9606 GN=ACTR3B PE=2 SV=1 |
| API5 | 73 | 59310 | 2 | 2 | 0.17 | Apoptosis inhibitor 5 OX=9606 GN=API5 PE=1 SV=3 |
| 1433B | 68 | 28179 | 2 | 2 | 0.4 | 14-3-3 protein beta/alpha OX=9606 GN=YWHAB PE=1 SV=3 |
| CC110 | 67 | 97235 | 2 | 1 | 0.05 | Coiled-coil domain-containing protein 110 OX=9606 GN=CCDC110 PE=1 SV=1 |

| | | | | | | |
|---|---|---|---|---|---|---|
| RL13A | 65 | 23619 | 2 | 2 | 0.49 | 60S ribosomal protein L13a OX=9606 GN=RPL13A PE=1 SV=2 |
| DHE4 | 65 | 61738 | 2 | 2 | 0.17 | Glutamate dehydrogenase 2, mitochondrial OX=9606 GN=GLUD2 PE=1 SV=2 |
| MTX2 | 62 | 30086 | 2 | 2 | 0.37 | Metaxin-2 OX=9606 GN=MTX2 PE=1 SV=1 |
| ABCD3 | 58 | 75941 | 2 | 2 | 0.13 | ATP-binding cassette sub-family D member 3 OX=9606 GN=ABCD3 PE=1 SV=1 |
| ARF1 | 56 | 20741 | 2 | 2 | 0.57 | ADP-ribosylation factor 1 OX=9606 GN=ARF1 PE=1 SV=2 |
| RAB10 | 52 | 22755 | 2 | 2 | 0.51 | Ras-related protein Rab-10 OX=9606 GN=RAB10 PE=1 SV=1 |
| LMNB1 | 45 | 66653 | 2 | 2 | 0.15 | Lamin-B1 OX=9606 GN=LMNB1 PE=1 SV=2 |
| RL36A | 44 | 12718 | 2 | 2 | 1.07 | 60S ribosomal protein L36a OX=9606 GN=RPL36A PE=1 SV=2 |
| PDIA1 | 41 | 57480 | 2 | 2 | 0.18 | Protein disulfide-isomerase OX=9606 GN=P4HB PE=1 SV=3 |
| DYH8 | 41 | 517984 | 2 | 1 | 0.01 | Dynein heavy chain 8, axonemal OX=9606 GN=DNAH8 PE=1 SV=2 |
| NEK10 | 40 | 134145 | 2 | 1 | 0.04 | Serine/threonine-protein kinase Nek10 OX=9606 GN=NEK10 PE=2 SV=3 |
| NUMA1 | 39 | 239199 | 2 | 2 | 0.04 | Nuclear mitotic apparatus protein 1 OX=9606 GN=NUMA1 PE=1 SV=2 |
| CC88B | 38 | 165166 | 2 | 1 | 0.03 | Coiled-coil domain-containing protein 88B OX=9606 GN=CCDC88B PE=1 SV=1 |
| MED16 | 37 | 98441 | 2 | 1 | 0.05 | Mediator of RNA polymerase II transcription subunit 16 OX=9606 GN=MED16 PE=1 SV=2 |
| CTBP2 | 31 | 49427 | 2 | 2 | 0.21 | C-terminal-binding protein 2 OX=9606 GN=CTBP2 PE=1 SV=1 |
| TNIK | 31 | 155361 | 2 | 2 | 0.06 | TRAF2 and NCK-interacting protein kinase OX=9606 GN=TNIK PE=1 SV=1 |
| SIPA1 | 31 | 112821 | 2 | 2 | 0.09 | Signal-induced proliferation-associated protein 1 OX=9606 GN=SIPA1 PE=1 SV=1 |
| CING | 30 | 136532 | 2 | 1 | 0.04 | Cingulin OX=9606 GN=CGN PE=1 SV=2 |
| CENPB | 30 | 65531 | 2 | 2 | 0.16 | Major centromere autoantigen B OX=9606 GN=CENPB PE=1 SV=2 |
| RASLC | 29 | 29872 | 2 | 1 | 0.17 | Ras-like protein family member 12 OX=9606 GN=RASL12 PE=1 SV=1 |
| SKAP1 | 26 | 41692 | 2 | 1 | 0.12 | Src kinase-associated phosphoprotein 1 OX=9606 GN=SKAP1 PE=1 SV=3 |
| TRI55 | 25 | 61454 | 2 | 1 | 0.08 | Tripartite motif-containing protein 55 OX=9606 GN=TRIM55 PE=1 SV=2 |
| SNPC4 | 25 | 160134 | 2 | 2 | 0.06 | snRNA-activating protein complex subunit 4 OX=9606 GN=SNAPC4 PE=1 SV=1 |
| ASPM | 23 | 413189 | 2 | 1 | 0.01 | Abnormal spindle-like microcephaly-associated protein OX=9606 GN=ASPM PE=1 SV=2 |
| KPRB | 23 | 41299 | 2 | 1 | 0.12 | Phosphoribosyl pyrophosphate synthase-associated protein 2 OX=9606 GN=PRPSAP2 PE=1 SV=1 |
| BCAR3 | 20 | 93476 | 2 | 1 | 0.05 | Breast cancer anti-estrogen resistance protein 3 OX=9606 GN=BCAR3 PE=1 SV=1 |
| MGAL | 17 | 279070 | 2 | 2 | 0.03 | Probable maltase-glucoamylase 2 OX=9606 GN=MGAM2 PE=2 SV=3 |
| NPM3 | 125 | 19617 | 1 | 1 | 0.27 | Nucleoplasmin-3 OX=9606 GN=NPM3 PE=1 SV=3 |
| MDHM | 119 | 35937 | 1 | 1 | 0.14 | Malate dehydrogenase, mitochondrial OX=9606 GN=MDH2 PE=1 SV=3 |
| IF3M | 88 | 31877 | 1 | 1 | 0.16 | Translation initiation factor IF-3, mitochondrial OX=9606 GN=MTIF3 PE=1 SV=4 |
| COPA | 86 | 139797 | 1 | 1 | 0.03 | Coatomer subunit alpha OX=9606 GN=COPA PE=1 SV=2 |
| HACD3 | 82 | 43360 | 1 | 1 | 0.11 | Very-long-chain (3R)-3-hydroxyacyl-CoA dehydratase 3 OX=9606 GN=HACD3 PE=1 SV=2 |
| VAPB | 79 | 27439 | 1 | 1 | 0.19 | Vesicle-associated membrane protein-associated protein B/C OX=9606 GN=VAPB PE=1 SV=3 |
| AP2S1 | 78 | 17178 | 1 | 1 | 0.31 | AP-2 complex subunit sigma OX=9606 GN=AP2S1 PE=1 SV=2 |
| MAP4 | 77 | 121443 | 1 | 1 | 0.04 | Microtubule-associated protein 4 OX=9606 GN=MAP4 PE=1 SV=3 |
| P5CS | 72 | 87989 | 1 | 1 | 0.06 | Delta-1-pyrroline-5-carboxylate synthase OX=9606 GN=ALDH18A1 PE=1 SV=2 |
| RS23 | 70 | 15969 | 1 | 1 | 0.34 | 40S ribosomal protein S23 OX=9606 GN=RPS23 PE=1 SV=3 |

### Supplemental Table 2 LetR only MS data – Proteins with ≥2 peptides or MASCOT score ≥67

| Accession | Score | Mass | No. of sig. matches | No. of sig. sequences | emPAI | Description |
|---|---|---|---|---|---|---|
| K1C15 | 3777 | 49409 | 137 | 9 | 1.6 | Keratin, type I cytoskeletal 15 OX=9606 GN=KRT15 PE=1 SV=3 |
| K2C6B | 2270 | 60315 | 136 | 13 | 1.99 | Keratin, type II cytoskeletal 6B OX=9606 GN=KRT6B PE=1 SV=5 |
| GFAP | 3583 | 49907 | 93 | 4 | 0.46 | Glial fibrillary acidic protein OX=9606 GN=GFAP PE=1 SV=1 |
| HSP72 | 1670 | 70263 | 34 | 12 | 1.24 | Heat shock-related 70 kDa protein 2 OX=9606 GN=HSPA2 PE=1 SV=1 |
| TBA3E | 388 | 50568 | 11 | 8 | 1.11 | Tubulin alpha-3E chain OX=9606 GN=TUBA3E PE=1 SV=2 |
| ADT3 | 302 | 33073 | 11 | 9 | 2.59 | ADP/ATP translocase 3 OX=9606 GN=SLC25A6 PE=1 SV=4 |
| STRBP | 216 | 74290 | 9 | 5 | 0.46 | Spermatid perinuclear RNA-binding protein OX=9606 GN=STRBP PE=1 SV=1 |
| TPM2 | 248 | 32945 | 7 | 4 | 0.77 | Tropomyosin beta chain OX=9606 GN=TPM2 PE=1 SV=1 |
| EF1A2 | 200 | 50780 | 7 | 6 | 0.75 | Elongation factor 1-alpha 2 OX=9606 GN=EEF1A2 PE=1 SV=1 |
| YBOX2 | 296 | 38552 | 6 | 4 | 0.63 | Y-box-binding protein 2 OX=9606 GN=YBX2 PE=1 SV=2 |
| H2A1D | 200 | 14099 | 6 | 5 | 4.2 | Histone H2A type 1-D OX=9606 GN=HIST1H2AD PE=1 SV=2 |
| IF2G | 188 | 51647 | 5 | 5 | 0.58 | Eukaryotic translation initiation factor 2 subunit 3 OX=9606 GN=EIF2S3 PE=1 SV=3 |
| NOLC1 | 138 | 73560 | 5 | 5 | 0.38 | Nucleolar and coiled-body phosphoprotein 1 OX=9606 GN=NOLC1 PE=1 SV=2 |
| RL26 | 94 | 17248 | 5 | 4 | 1.96 | 60S ribosomal protein L26 OX=9606 GN=RPL26 PE=1 SV=1 |
| MTMRE | 58 | 73013 | 5 | 1 | 0.07 | Myotubularin-related protein 14 OX=9606 GN=MTMR14 PE=1 SV=2 |
| ODO1 | 94 | 117059 | 4 | 4 | 0.18 | 2-oxoglutarate dehydrogenase, mitochondrial OX=9606 GN=OGDH PE=1 SV=3 |
| FLOT2 | 74 | 47434 | 4 | 4 | 0.49 | Flotillin-2 OX=9606 GN=FLOT2 PE=1 SV=2 |
| H2AV | 134 | 13501 | 3 | 2 | 0.99 | Histone H2A.V OX=9606 GN=H2AFV PE=1 SV=3 |
| EXOS2 | 52 | 32996 | 3 | 2 | 0.33 | Exosome complex component RRP4 OX=9606 GN=EXOSC2 PE=1 SV=2 |
| P5CR2 | 135 | 33958 | 2 | 2 | 0.32 | Pyrroline-5-carboxylate reductase 2 OX=9606 GN=PYCR2 PE=1 SV=1 |
| HNRH2 | 96 | 49517 | 2 | 2 | 0.21 | Heterogeneous nuclear ribonucleoprotein H2 OX=9606 GN=HNRNPH2 PE=1 SV=1 |
| DHB4 | 94 | 80092 | 2 | 2 | 0.13 | Peroxisomal multifunctional enzyme type 2 OX=9606 GN=HSD17B4 PE=1 SV=3 |
| SEPT7 | 84 | 50933 | 2 | 2 | 0.2 | Septin-7 OX=9606 GN=SEPT7 PE=1 SV=2 |
| ISOC2 | 76 | 22608 | 2 | 2 | 0.51 | Isochorismatase domain-containing protein 2 OX=9606 GN=ISOC2 PE=1 SV=1 |
| SEP11 | 64 | 49652 | 2 | 2 | 0.21 | Septin-11 OX=9606 GN=SEPT11 PE=1 SV=3 |
| MYH11 | 63 | 228054 | 2 | 2 | 0.04 | Myosin-11 OX=9606 GN=MYH11 PE=1 SV=3 |
| CALU | 62 | 37198 | 2 | 2 | 0.29 | Calumenin OX=9606 GN=CALU PE=1 SV=2 |
| IF4E2 | 61 | 28458 | 2 | 2 | 0.39 | Eukaryotic translation initiation factor 4E type 2 OX=9606 GN=EIF4E2 PE=1 SV=1 |
| 2AAA | 60 | 66065 | 2 | 2 | 0.15 | Serine/threonine-protein phosphatase 2A 65 kDa regulatory subunit A alpha isoform OX=9606 GN=PPP2R1A PE=1 SV=4 |
| SLIRP | 58 | 12398 | 2 | 2 | 1.11 | SRA stem-loop-interacting RNA-binding protein, mitochondrial OX=9606 GN=SLIRP PE=1 SV=1 |
| UBP47 | 56 | 158581 | 2 | 1 | 0.03 | Ubiquitin carboxyl-terminal hydrolase 47 OX=9606 GN=USP47 PE=1 SV=3 |
| TOP1M | 55 | 70398 | 2 | 2 | 0.14 | DNA topoisomerase I, mitochondrial OX=9606 GN=TOP1MT PE=1 SV=1 |
| RL22L | 54 | 14598 | 2 | 2 | 0.89 | 60S ribosomal protein L22-like 1 OX=9606 GN=RPL22L1 PE=1 SV=2 |
| CTNB1 | 49 | 86069 | 2 | 2 | 0.12 | Catenin beta-1 OX=9606 GN=CTNNB1 PE=1 SV=1 |
| TXND5 | 48 | 48283 | 2 | 2 | 0.22 | Thioredoxin domain-containing protein 5 OX=9606 GN=TXNDC5 PE=1 SV=2 |
| RAB43 | 48 | 23553 | 2 | 2 | 0.49 | Ras-related protein Rab-43 OX=9606 GN=RAB43 PE=1 SV=1 |
| SPTN2 | 45 | 272526 | 2 | 2 | 0.04 | Spectrin beta chain, non-erythrocytic 2 OX=9606 GN=SPTBN2 PE=1 SV=3 |
| COX6C | 45 | 8776 | 2 | 2 | 1.85 | Cytochrome c oxidase subunit 6C OX=9606 GN=COX6C PE=1 SV=2 |
| SRSF7 | 45 | 27578 | 2 | 2 | 0.41 | Serine/arginine-rich splicing factor 7 OX=9606 GN=SRSF7 PE=1 SV=1 |
| ATP5H | 43 | 18537 | 2 | 2 | 0.66 | ATP synthase subunit d, mitochondrial OX=9606 GN=ATP5H PE=1 SV=3 |
| AGR2 | 41 | 20024 | 2 | 2 | 0.6 | Anterior gradient protein 2 homolog OX=9606 GN=AGR2 PE=1 SV=1 |

| | | | | | | |
|---|---|---|---|---|---|---|
| GOGB1 | 41 | 377215 | 2 | 1 | 0.01 | Golgin subfamily B member 1 OX=9606 GN=GOLGB1 PE=1 SV=2 |
| EIF3M | 41 | 42932 | 2 | 2 | 0.25 | Eukaryotic translation initiation factor 3 subunit M OX=9606 GN=EIF3M PE=1 SV=1 |
| AP5B1 | 32 | 95087 | 2 | 1 | 0.05 | AP-5 complex subunit beta-1 OX=9606 GN=AP5B1 PE=1 SV=4 |
| RING2 | 32 | 38088 | 2 | 2 | 0.28 | E3 ubiquitin-protein ligase RING2 OX=9606 GN=RNF2 PE=1 SV=1 |
| PLSI | 32 | 70608 | 2 | 2 | 0.14 | Plastin-1 OX=9606 GN=PLS1 PE=1 SV=2 |
| FA5 | 31 | 252686 | 2 | 2 | 0.04 | Coagulation factor V OX=9606 GN=F5 PE=1 SV=4 |
| PAB4L | 29 | 42056 | 2 | 2 | 0.25 | Polyadenylate-binding protein 4-like OX=9606 GN=PABPC4L PE=2 SV=1 |
| EXOS7 | 28 | 32428 | 2 | 2 | 0.34 | Exosome complex component RRP42 OX=9606 GN=EXOSC7 PE=1 SV=3 |
| NOP10 | 25 | 7758 | 2 | 1 | 0.8 | H/ACA ribonucleoprotein complex subunit 3 OX=9606 GN=NOP10 PE=1 SV=1 |
| NPIL2 | 20 | 46117 | 2 | 1 | 0.11 | Putative NPIP-like protein LOC729978 OX=9606 PE=5 SV=4 |
| NHRF2 | 80 | 37619 | 1 | 1 | 0.13 | Na(+)/H(+) exchange regulatory cofactor NHE-RF2 OX=9606 GN=SLC9A3R2 PE=1 SV=2 |
| RUXGL | 73 | 8595 | 1 | 1 | 0.71 | Putative small nuclear ribonucleoprotein G-like protein 15 OX=9606 GN=SNRPGP15 PE=5 SV=2 |
| IBP5 | 67 | 31576 | 1 | 1 | 0.16 | Insulin-like growth factor-binding protein 5 OX=9606 GN=IGFBP5 PE=1 SV=1 |

| **Overlap over known AR interactors** | |
|---|---|
| ACTB | ACL6A |
| ACTN4 | ANDR |
| CALM1 | APEX1 |
| CCNH | ATPB |
| DDX17 | ATPO |
| DDX5 | ATX2L |
| FLNA | CSTF1 |
| FOXA2 | CSTF3 |
| HDAC1 | CTBP1 |
| HDAC7 | CTBP2 |
| HMGB1 | DDX42 |
| KAT2A | DDX53 |
| KDM1A | DLDH |
| KDM3A | ETFB |
| MCM7 | EXOS2 |
| MED24 | FA98B |
| MYO6 | FUBP3 |
| NONO | HNRL1 |
| NSD1 | KDM1A |
| PA2G4 | LSM12 |
| PAWR | MTA2 |
| PHB | NOLC1 |
| PPID | ODO1 |
| PRDX1 | ODPA |
| PRKDC | ODPB |
| PSPC1 | PP1B |
| RAN | PRPF3 |
| RBM14 | RBBP4 |
| RREB1 | RBM10 |
| SART3 | RL18A |
| SFPQ | RL34 |
| SGTA | RLA1 |
| SMAD3 | RS11 |
| SUMO2 | RS30 |
| XRCC5 | RTRAF |
| XRCC6 | SMCA2 |
| XRN2 | SMCE1 |
| | SMD2 |
| | SNRPA |
| | U2AF1 |

Supplemental Table 3 AR interacting proteins present in Mass Spectrometry analysis also present in published AR RIME data and AR interacting proteins and coregulators table.
